# Marine community metabolomes in the eastern tropical North Pacific Oxygen Deficient Zone reveal glycine betaine as a metabolic link between *Prochlorococcus* and SAR11

**DOI:** 10.1101/2025.05.07.652674

**Authors:** Natalie A. Kellogg, Clara A. Fuchsman, Laura T. Carlson, Robert M. Morris, Anitra E. Ingalls, Gabrielle Rocap

## Abstract

Oxygen deficient zones (ODZs) are subsurface marine systems that harbor distinct microbial communities, including populations of the picocyanobacteria *Prochlorococcus* that can form a secondary chlorophyll maximum (SCM), and low-oxygen tolerant strains of the globally abundant heterotroph *Pelagibacter* (SAR11). Yet, the small labile molecules (metabolites) responsible for maintaining these ODZ communities are unknown. Here, we compared the metabolome of an ODZ to that of an oxygenated site by quantifying 87 metabolites across depth profiles in the eastern tropical North Pacific ODZ and the oxygenated waters of the North Pacific Gyre. Metabolomes were largely consistent between anoxic and oxic water columns. However, the osmolyte glycine betaine (GBT) was enriched in the oxycline and SCM of the ETNP, comprising as much as 1.2% of particulate organic carbon. Transcriptomes revealed two active GBT production pathways, glycine methylation (*SDMT/bsmB*) expressed by *Prochlorococcus* and choline oxidation (*betB*) expressed by Gammaproteobacteria. GBT consumption through demethylation involved diverse microbial taxa, with SAR11 contributing nearly half of the transcripts for the initial step of GBT demethylation (BHMT), which is predicted to convert GBT and homocysteine into dimethylglycine and methionine, a compound SAR11 cannot otherwise produce. Thus, GBT connects the metabolisms of the dominant phototroph and heterotroph in the oceans.

## Introduction

Annually, marine phytoplankton fix up to 50 billion tons of carbon (Falkowski, 1994), with roughly half of this carbon being metabolized by marine bacteria (Azam et al., 1983). Freshly synthesized small biomolecules, known as metabolites, are among the most readily used compounds, playing an important role in fostering taxa interdependencies that structure microbial communities (Poretsky et al., 2010; Vorobev et al., 2018; Boysen et al., 2022; Moran et al., 2022). Relatively few studies have quantified changes in metabolite pools across vertical gradients in the marine environment (Johnson et al., 2020; Heal et al., 2021; Johnson et al., 2023; Longnecker et al., 2024), particularly with respect to oxygen. Oxygen deficient zones (ODZs) are permanent naturally occurring subsurface regions bounded by steep clines in oxygen concentrations between oxygen-rich surface and deep waters above and below the completely anoxic midwater layer, where distinct microbial communities thrive and mediate elemental cycles (Beman and Carolan, 2013; Ganesh et al., 2014; Fuchsman et al., 2017; Wakeham, 2020). Here we compare metabolomes of natural communities in regions of the North Pacific Ocean with and without a subsurface ODZ to investigate whether metabolites produced by the distinct communities within ODZs differ from those in neighboring aerobic waters. We further use transcriptomes to identify taxa involved in production and subsequent transformation of glycine betaine (GBT), a metabolite we suggest is involved in microbial interdependencies in this community, and elsewhere in the ocean.

Secondary chlorophyll maxima (SCM) are often observed in ODZs when the top of the anoxic layer is shallow enough to intersect with the bottom of the euphotic zone. In these waters nutrient concentrations are relatively high (Fuchsman et al., 2017; Wakeham, 2020), light levels are very low (1% blue light) (Cepeda-Morales et al., 2009) and the photoautotrophic community in the SCM is almost entirely composed of the unicellular cyanobacteria *Prochlorococcus* (Johnson et al., 1999; Goericke et al., 2000). Multiple genetically distinct lineages (ecotypes) of *Prochlorococcus* coexist within the ODZ SCM, including LL IV, AMZ Ia and Ib (formerly LL V), AMZ II (formerly LL VI) and AMZ III (Lavin et al., 2010; Fuchsman et al., 2019; Ulloa et al., 2021). With the exception of LL IV, these ecotypes have not been detected outside of ODZs and to date none of the AMZ ecotypes have been cultured. Metagenomic and single-cell amplified genome (SAG) analyses suggest that AMZ *Prochlorococcus* ecotypes may be specifically adapted in different ways to the environmental conditions within ODZs. They have the gene repertoire for ammonium, urea, nitrite, and nitrate use and some have the genetic potential to produce complete phycobilisomes (Widner et al., 2018; Ulloa et al., 2021), while others contain a second form II Rubsico (Jaffe et al., 2024). They also possess genes that may provide adaptations to their low oxygen environment of the ODZ, including for nitric oxide detoxification, alcohol fermentation, and oxygen-independent tetrapyrrole biosynthesis (Ulloa et al., 2021). The O_2_ and organic matter that these *Prochlorococcus* produce are presumably consumed by the surrounding microbial community in ODZs, potentially driving the production of N_2_O—a potent greenhouse gas—by ammonia-oxidizing Thaumarcheota and N_2_ production by denitrifying bacteria, contributing to nitrogen loss from the ocean (Garcia-Robledo et al., 2017; Fuchsman et al., 2019). It is estimated that *Prochlorococcus* provides up to 40% of the organic matter used by the heterotrophic community in the SCM (Fuchsman et al., 2019).

Members of the ubiquitous Pelagibacterales (SAR11) are also abundant in ODZs, including in anoxic waters where they can account for 30-40% of the microbial community (Tsementzi et al., 2016; Fuchsman et al., 2017; Ruiz-Perez et al., 2021). SAR11 bacteria are the most abundant lineage of bacteria in the ocean (Morris et al., 2002) and are major consumers of dissolved organic matter produced by phytoplankton (Giovannoni, 2017). Like *Prochlorococcus*, this heterotrophic lineage is also comprised of diverse subclades, including ecotypes specific to ODZs (clades 1c and IIa.A) that remain uncultured but have the genomic potential to reduce nitrate to nitrite via a respiratory nitrate reductase, the first step of denitrification (Tsementzi et al., 2016; Ruiz-Perez et al., 2021).The SAR11 group as a whole have streamlined genomes that suggest the loss of key biosynthetic genes and thus reliance on other community members to provide specific biomolecules for growth (Giovannoni 2017). They lack Vitamin B12 biosynthesis genes as well as both the Vitamin B12-independent *MetE* and -dependent *MetH* pathways for methionine synthesis. Cultured strains have specific growth requirements for glycine and pyruvate (or precursors) and organosulfur compounds (Carini et al., 2012; Giovannoni, 2017). In co-culture, a strain of LL IV *Prochlorococcus* capable of synthesizing GBT was able to meet the glycine requirement of the surface-dwelling Ia.3 ecotype SAR11 strain, while a strain of HL I *Prochlorococcus* lacking genes for GBT synthesis could not (Becker et al., 2019).

Here, we measure the in-situ inventory of small polar metabolites in the particulate (presumably intracellular) pool within depth profiles at three stations to facilitate a deeper understanding of marine microbial physiology and to identify the interactions that drive ecosystem diversity and activity in ODZs. Intracellular metabolite profiles of model marine microbes are taxon-specific and can respond to environmental perturbations (Heal et al., 2019; Boysen et al., 2021; Heal et al., 2021; Johnson et al., 2020; Durham et al., 2022; Johnson et al., 2023; Kujawinski et al., 2023). However, measurements of the chemical diversity and concentrations of metabolites present in ODZs remain unknown. We quantified 87 metabolites along depth profiles in the tropical North Pacific Gyre (NPG) and ETNP ODZ. We emphasize the ubiquitous osmolyte glycine betaine (GBT), a major compatible solute produced by select heterotrophic bacteria and phytoplankton (Boysen et al., 2022), including the picocyanobacteria *Synechococcus* and some deeply-branching *Prochlorococcus* lineages (Klähn et al., 2010; Klähn and Hagemann, 2011; Kujawinski et al., 2023). GBT has the potential to perform various biochemical functions within the ODZ community, which may be taxon-specific, including as a source of energy, carbon, nitrogen, other metabolites like glycine, or as a methyl group donor (Boysen et al., 2022). Using metatranscriptomes, we taxonomically resolve the metabolic pathways for GBT synthesis and transformation. We show that within the oxycline and SCM of the ETNP, *Prochlorococcus* and SAR11 are direct producers and consumers, respectively, of GBT. The ability to use GBT as a methyl donor in methionine synthesis and as a glycine precursor is likely key to the survival of SAR11 in the ODZ, and elsewhere in the oceans.

## Materials and Methods

### Sample collection for metabolomic analysis

Samples were collected in the North Pacific for metabolomics analysis of particulate material within the upper 3000 m (**Figure 1, Table S1**). Samples for the oxic North Pacific gyre (NPG) depth profile were collected during MGL1704 (Gradients II) aboard the R/V *Marcus Langseth* at eight depths between 15 and 3000 m on June 10, 2017 (St 17; 32°N 157.6°W). Additional depth profiles from a more Northern transition zone station and a more Southern gyre station from this same cruise are also available (Heal et al., 2021). During RR1805 (POMZ 2018), samples for the eastern tropical North Pacific (ETNP) ODZ depth profiles were taken at two stations, one coastal (St. P1; 20.3 °N 106.1°W) at eight depths between 10 and 1500 m and one offshore (St. P2; 16.9°N 107°W) at eight depths between 10 and 400 m, from April 14, 2018 to May 2, 2018 aboard the R/V *Roger Revelle*. At each sampling location, single or triplicate filters were collected for environmental metabolomics, as described in Boysen et al. 2018, using 10-L Niskin bottles on a CTD-rosette. Each sample set was collected between 1 and 4 PM. At each depth, 10 L of seawater was collected into polycarbonate carboys and the seawater particulates were harvested by gentle filtration onto 142 mm 0.2-µm PTFE Omnipore Membrane filters using a peristaltic pump, flash frozen in liquid N_2_, and stored at −80°C. Methodological blanks were collected in triplicate filters by pre-filtering through one 0.2µm PTFE Omnipore Membrane filters and passing the filtrate though a clean filter. This blank was used to remove contaminants and compounds adsorbed onto the filter from the dissolved pool.

**Figure 1:**
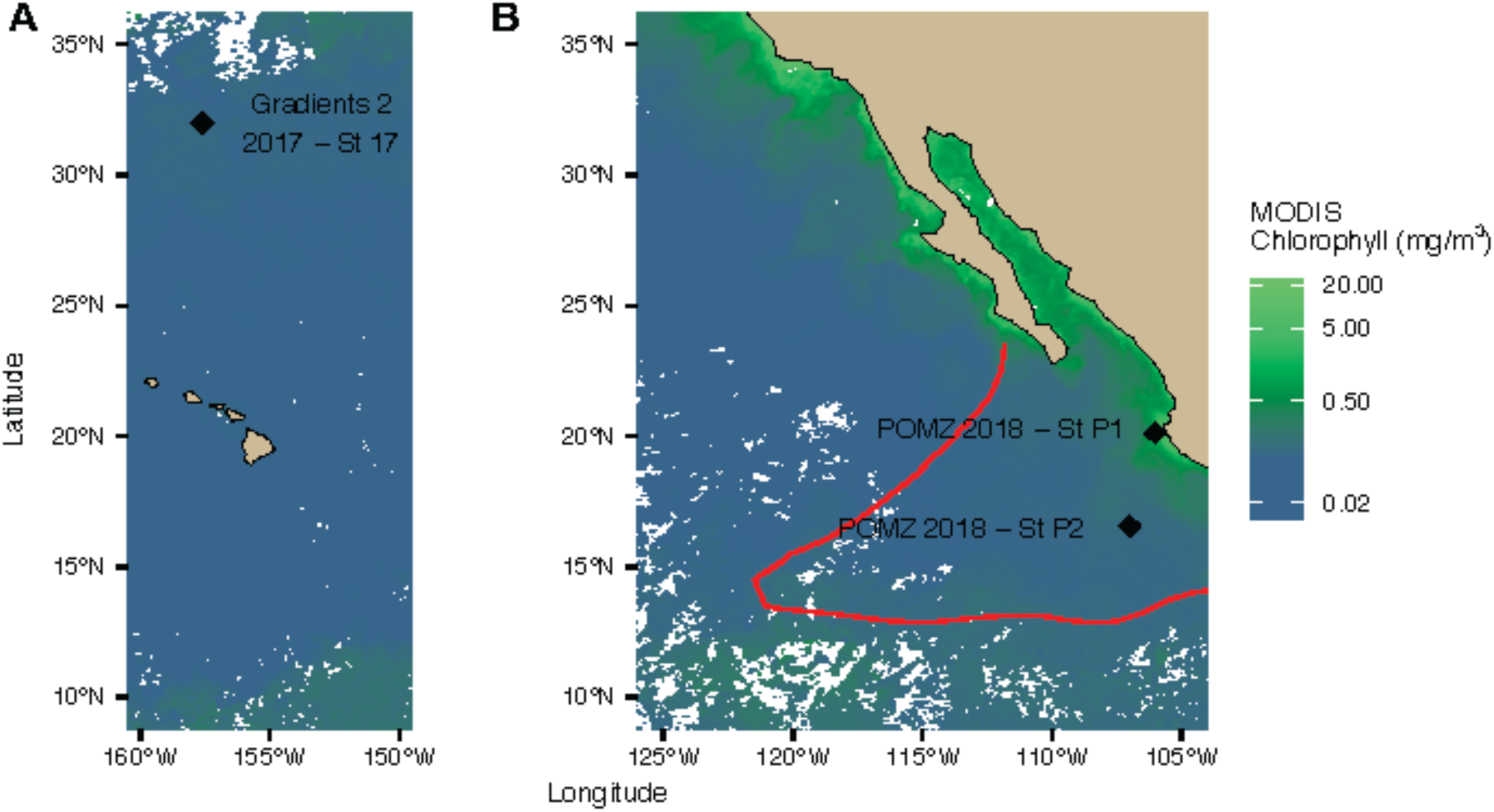
Map of stations sampled. MODIS satellite chlorophyll is from the time of sampling during **A)** MGL1704 (Gradients 2 2017; Station 17, ∼0.07 mg/m^3^ chlorophyll) and **B)** RR1805 (POMZ 2018; Stations P2, ∼0.11 mg/m³ chlorophyll, and P1, ∼1.27 mg/m³ chlorophyll). Red line indicates the 10 µM oxygen deficient zone (ODZ) boundary at 300 meters as determined from World Ocean Atlas (Boyer et al., 2018).

### Metabolite extraction

Metabolites were extracted with a modified Bligh and Dyer method (Bligh and Dyer, 1959; Boysen et al., 2018) using a solvent mixture of methanol, water, and dichloromethane in a 1:1:2 ratio and bead-beaten three times for 30 seconds each over a 30-minute period, with samples kept at −20°C between intervals. This process separated the sample into two distinct fractions: a polar aqueous extract (comprising methanol- and water-soluble components) and an organic extract (containing dichloromethane-soluble components). The sample extracts were dried under clean N_2_ and reconstituted in 400 µl water. Only the metabolites in the aqueous phase were analyzed for this study.

### Liquid chromatography-mass spectrometry

Metabolomics analysis was done using liquid chromatography-mass spectrometry (LC-MS) as reported in Boysen et al., 2018. Both reversed phase (RP - Waters Acquity UPLC HSS Cyano column, 1.8 *µ*m particle size, 2.1 mm x 5 mm) and hydrophilic interaction (HILIC - SeQuant ZIC-pHILIC column, 5 mm particle size, 2.1 mm x 150 mm, from Millipore) chromatography were used to analyze the non-polar and polar compounds in the aqueous phase extract, respectively. Internal standards were then added at the same concentrations as in (Boysen et al., 2018) to use in normalization and quantification. Both LC columns were connected to a Thermo QExactive HF Orbitrap (QE) mass spectrometer for high resolution, accurate mass analysis (Boysen et al. 2018). For HILIC, a full scan method with polarity switching was used at a resolution of 60,000. For RP, positive ionization mode was used with a resolution of 120,000. Raw data files (.raw) were converted to .mzXML format using Proteowizard (Kessner et al., 2008) for downstream analysis and deposited to Metabolomics Workbench (Sud et al., 2016).

### Metabolite data processing

Chromatographic peaks were integrated in Skyline (MacLean et al. 2010) and subjected to in-house quality control. Peaks that did not meet minimum quality criteria (signal:noise ratio, minimum peak area, comparison to methodological blank area, retention time, mass error) were excluded from further analysis and replaced with zeros for statistical analysis. If peak areas passed quality control they were normalized using the best-match internal standard (B-MIS) package (Boysen et al., 2018) to control for method and instrument variability. Details are provided in Text S1.

### Metabolite concentration calculations

Absolute concentrations were calculated for 87 identified peaks using commercially available standards run in the same batch as the environmental samples, in a similar method as described in (Heal et al., 2021). For metabolites where isotopically labeled internal standards were spiked into the samples, concentrations were calculated using isotope-labeled peak areas. For compounds without internal standards, a response factor (RF) and RF ratio was calculated for each compound to correct for ion suppression using authentic standards mixed into water and a representative matrix (**Text S1**). Particulate metabolite concentrations were then normalized to volume of water filtered resulting in a nanomolar environmental concentration (nmol L^−1^). In some instances, metabolite concentrations (nM compound) are converted to carbon or nitrogen units (nM C or nM N in a metabolite) by multiplying the nmols of a metabolite by the number of carbon or nitrogen atoms in each metabolite (**Table S2, S3**).

### Multivariate statistics

For multivariate analyses, a data matrix of 87 quantified compounds for each biological replicate were obtained and standardized to the maximum value of each metabolite across samples. Differences in sample types, grouped *a priori* by measured environmental characteristics, were analyzed using a non-metric multidimensional scaling (NMDS) approach with Euclidian distance. The surface is defined as the upper water column, where fluorescence remains relatively constant. The deep chlorophyll maximum (DCM) represents the highest fluorescence point near the surface. The oxycline is described as a sharp drop in oxygen from >200 µmol kg^−1^ in the surface to zero, while the SCM corresponds to the point where peak fluorescence occurs below the DCM, with no detectable oxygen. The mid-anoxic and mid-oxic zones extend from 120 to 600 meters, with the mid-anoxic zone lacking oxygen and the mid-oxic zone containing measurable levels. Deep oxic samples were taken from depths of 1,000 meters and beyond, where oxygen is present (**Table S1**). The dimensionality of the NMDS was evaluated using a scree plot, and a 2-dimensional NMDS solution was selected, with the probability assessed through a Monte Carlo permutation. An analysis of similarities (ANOSIM) was performed on environmentally defined groups across the depth profile samples, as well as between stations, to determine whether differences existed in the particulate metabolite data between the *a priori* defined groups or between stations. Indicator species analysis was then conducted using the same environmentally defined groupings to explore the relationships between metabolite abundance and sample types (**Table S4**). Data transformation, normalization, NMDS, double dendrogram heatmaps, ANOSIMS, and indicator species analysis statistics were performed using the R packages vegan (v2.5-7), pheatmap (v1.0.12), or indicspecies (v1.7.8).

### Metatranscriptomics

Previously published metatranscriptomes from the ETNP in the upper oxycline and the top of the ODZ (oxygen range: 0-33 µM; depth range 28-150 m) were collected from the same stations as the metabolite data on April 14, 2018 to May 2, 2018 aboard the R/V *Roger Revelle* and processed as previously described (Mattes et al., 2022). Briefly, between 3.5 to 4.2 L seawater was vacuum-filtered onto 0.2 μm Isopore membrane filters (Millipore, Inc., Billerica, MA) under a nitrogen headspace that limited oxygen diffusion during filtration. RNA was extracted with the Direct-zol RNA mini-prep plus kit (Zymo Research Corporation, Irvine, CA) and further subjected to DNase I treatment using the TURBO DNA-free kit (Thermo Fisher Scientific, Waltham, MA) and rRNA depletion with the Ribo-Zero rRNA removal kit for bacteria (Illumina, Inc., San Diego, CA). To generate quantitative transcript inventories, a set of 14 internal mRNA standards synthesized as previously described (Satinsky et al., 2013) were also added to the extraction buffer. Sequencing libraries were prepared with the Illumina TruSeq library prep kit v2 for paired-end (2 × 150 bp) sequencing and sequenced in four technical replicates each on an Illumina MiSeq instrument at the University of Washington Northwest Genomics Center (Seattle, WA, USA) which yielded between 20,038,590 and 30,533,274 raw reads per sample. Sequences were quality controlled using MG-RAST v 4.0.3 (Meyer et al. 2008) to trim adapters, filter for length and sequence quality, remove artificial duplicate reads and screen for human sequence contamination (MG-Rast project: ETNP ODZ Metatranscriptomes, mgp92168). Technical replicates from each sample were downloaded from MG-RAST after these quality control steps and combined. Reads were then further filtered to remove remaining ribosomal RNA sequences (between 1-8% of remaining reads) using SortMeRNA (version 2.1b) (Kopylova et al., 2012) and normalized to internal standards as previously described (Satinsky et al., 2013) (Table **S5**).

ETNP metagenomic reads from 60-300 m at St. P2 in 2012 (Fuchsman et al 2017) were assembled together in a combined assembly using MEGAHIT v1.12 (Li et al., 2015) in paired end mode with a minimum contig length of 500 (GenBank accession: GCA_002998485.1). Genes were identified using Prodigal v2.6.3 (Hyatt et al., 2010). Taxonomic assignments used kaiju (Menzel et al., 2016) and a custom database of MarDB v 1.5, MarRef v 6, MMETSP, and NCBI viral RefSeq v 208 (Keeling et al., 2014; Brister et al., 2015; O’Leary et al., 2016; Klemetsen et al., 2018). Functional annotations were done with KofamScan (Aramaki et al., 2020) using KEGG v113 profiles downloaded on February 12, 2025.

Genes associated with glycine betaine (GBT) synthesis and transformation were identified based on KEGG orthologs. A full list of GBT-related KEGG pathways and orthologs used in this analysis is provided in **Table S6**, along with metagenome gene presence and transcript abundance data summarized across depth and station (**Table S7**). Metatranscriptomic reads were mapped to annotated genes from the co-assembled metagenome using Bowtie2 v2.4.5 (Langmead and Salzberg, 2012), and gene-specific read counts were used to quantify pathway-level expression.

### Phylogenetic analysis

To assign taxonomy of *Prochlorococcus* ecotypes to expression of sarcosine dimethylglycine methyltransferase (*SDMT*/*bsmB*), a reference phylogenetic tree was constructed using both assembled proteins from the ETNP 2012 metagenomes, ETSP cyanobacterial SAGs (Ulloa et al., 2021), and other cyanobacterial sequences taken from MARDB v 1.5 (Klemetsen et al., 2018). Custom BLAST databases (Altschul et al., 1997) were created using proteins from the assembled ETNP contigs and proteins from single cell cyanobacterial genomes. Appropriate genes were obtained by blasting known reference sequences (blastp) against this database. All sequences recruited from blast were combined with the previously published full-length gene sequences and aligned in amino acid space with MUSCLE v.4 (Edgar, 2004). Maximum likelihood phylogenetic trees were constructed using the program IQ-TREE v. 2.2.5 (Nguyen et al., 2015). The tree was constructed with a gamma model of rate heterogeneity, and the amino acid substitution model Q.pfam+R5 was used. Bootstrap analyses (n = 100) were performed on the tree.

### Oceanographic data

Additional ancillary measurements were collected to characterize the environmental and biological setting. Satellite chlorophyll concentrations were taken from MODIS-Aqua at 8-day, 9-km resolution over the time periods of profile sampling. For both cruises, a Seabird 911 Conductivity Temperature Density meter, a Seabird SBE 43 Dissolved Oxygen Sensor, a WETLabs ECO Chlorophyll Fluorometer, and a Biospherical/Licor PAR/Irradiance Sensor were attached to the rosette. CTD and nutrient data for the R/V *Revelle* are available at BCO-DMO (dataset 779185). Nutrient samples were prefiltered with 0.2 µm Sterivex filters and analyzed at the UW Marine Chemistry Lab using a Technicon AAII system as described by the World Ocean Circulation Experiment (WOCE) Hydrographic Program protocol (Gordon et al., 1993). For ETNP particulate organic carbon samples from St. P2 and St. P1, 4 L of water were obtained from Niskin bottles and filtered onto pre-combusted GF/F filters as described in Fuchsman et al., 2025. Samples were fumed with HCl overnight to remove carbonate, dried at 40°C, packed in both silver and tin capsules, and sent to the UC Davis Stable Isotope Facility for analysis utilizing an Elemental Analyzer (Elementar Vario EL Cube) attached to an Isotope Ratio Mass Spectrometer (Isoprim VisION). Blank combusted GF/F filters did not show measurable signal. Particulate C from the NPG was collected on pre-combusted GF/F filters, processed following HOT protocols and combusted using an Exeter Analytical CE-440 CHN elemental analyzer (https://hahana.soest.hawaii.edu/hot/methods/pcpn.html). This protocol does not remove carbonates but suspended particulate inorganic C (carbonates) are at very low concentrations in the NPG and can be considered negligible (Karl et al., 2022).

## Results

### Environmental context

The water column structure of the sampled sites demonstrated three distinct environments within the tropical North Pacific Ocean: one oxic subtropical gyre profile and two tropical ODZ profiles, offshore and coastal (**Figure 1**). The subtropical NPG station chlorophyll fluorescence profile had a pronounced subsurface peak at 90 m, indicating the location of the deep chlorophyll maximum (DCM), with ambient oxygen concentrations of ∼240 µmol kg^−1^ (**Figure 2A**). This station provides a useful comparison to offshore (St. P2) and coastal (St. P1) ODZ stations, where oxygen concentrations dropped from >200 µmol kg^−1^ in the surface layer to below detection within the core of the ODZ.

**Figure 2:**
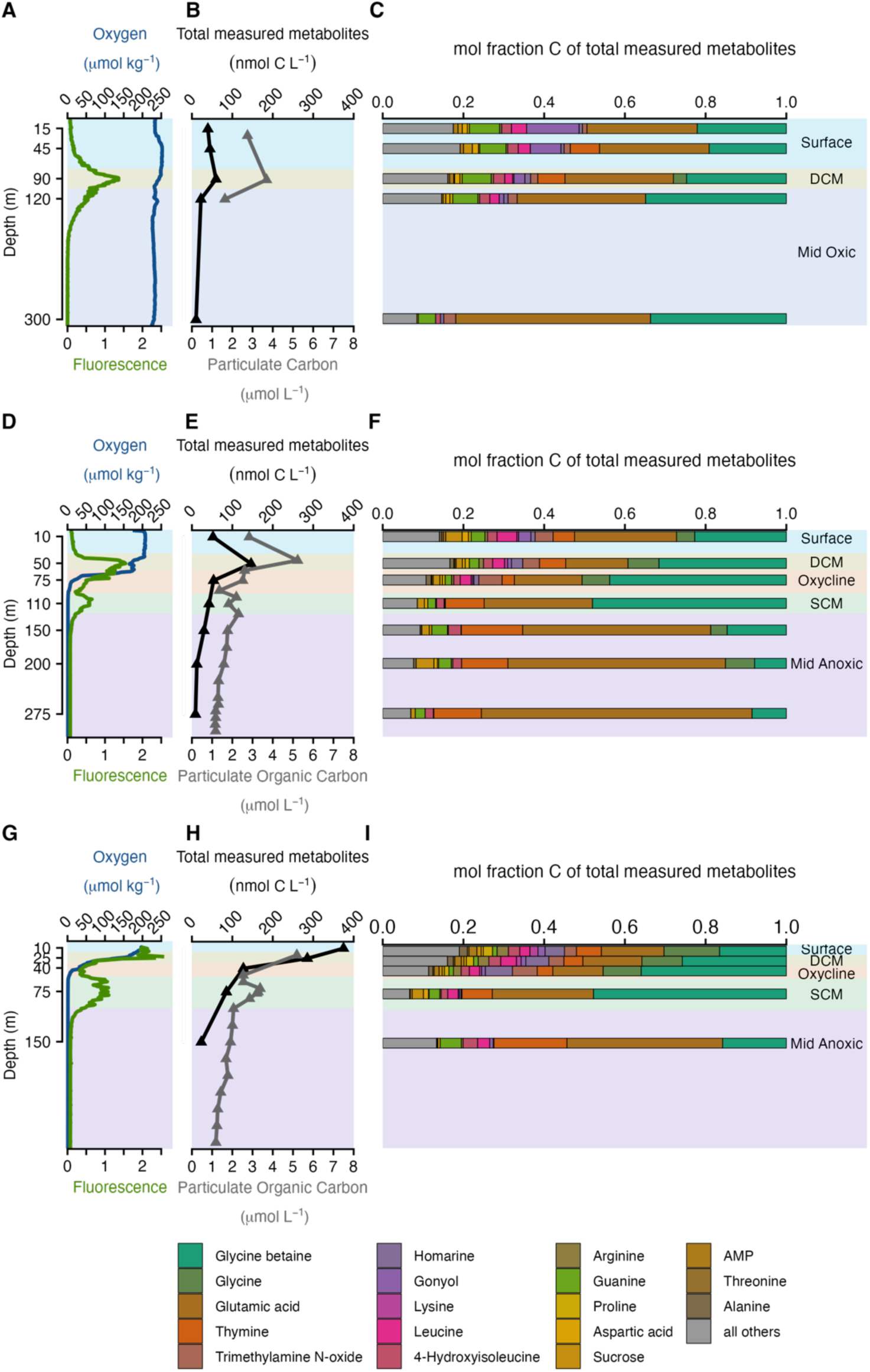
Most abundant 18 metabolites in environmental samples, presented as mole fraction of carbon of total identified metabolites, paired with total measured metabolites, particulate carbon (PC) and particulate organic carbon (POC), and CTD measurements of oxygen and fluorescence. Depth profiles from oxic NPG **(A-C)**, offshore ODZ (St. P2) **(D-F)**, and coastal ODZ (St. P1) **(G-I)**. Locations of samples are shown in Figure 1.

At both offshore and coastal ODZ stations **(Figure 1)** the chlorophyll fluorescence profiles featured two distinct subsurface peaks, the DCM and the SCM **(Figure 2D, 2G)**. Based on the Seabird SBE 43 oxygen sensor, at both of these stations, the DCM was located in oxic waters while the SCM occurred just below the oxycline at the top of the ODZ. The offshore ODZ station had a deeper DCM (50 m), oxycline (60-90 m), and SCM (110 m) compared to the coastal ODZ station’s DCM (25 m), oxycline (25-50m), and SCM (75 m) **(Figure 2D, 2G)**. While the oxygen sensor used on the 2018 cruise had a detection limit of ∼1 µM, these exact stations were previously shown to be functionally anoxic (<10 nM oxygen) using a STOX sensor (Tiano et al., 2014; Fuchsman et al., 2019). Although a STOX sensor was not available in the 2018 cruise analyzed here, nitrite accumulation above 0.5 µM is considered an indicator of anoxia (Banse et al., 2017). Nitrite concentrations were elevated at St. P2 between 110 and 500 m, reaching 3 µM at 140 m, and similarly elevated at St. P1 from 55 to 700 m, with concentrations reaching 3.6 µM at 125 m **(Figure S1)**, indicating that these layers were anoxic.

### Metabolite abundance and patterns

Metabolite concentrations were generally higher at the ODZ stations compared to the oxic NPG station. Across all profiles, absolute concentrations of 87 metabolites were determined (**Table S2**), ranging from 4.2 to 374.9 nM particulate carbon (**Figure 2B, 2E, 2H, Table S3**). These concentrations tracked closely with particulate carbon profiles, peaking in the oxygenated surface or DCM layers and declining with depth (Figure 2B, 2E, 2H).In the oxic NPG profile, metabolites ranged from 4.2 to 59.6 nM carbon, representing roughly ∼1.6% of total particulate carbon in the oxic DCM sample (**Figure 2B**). In the offshore ODZ, metabolite concentrations of 4.3 to 146.4 nM carbon made up ∼2.8% of total particulate organic carbon in the oxic DCM sample and ∼1.5% in the anoxic SCM sample (**Figure 2E**). The coastal ODZ profile had the highest metabolite concentrations, ranging from 7.1 to 374.9 nM carbon, representing 5.5% and 2.6% of particulate organic carbon in oxic DCM and anoxic SCM sample, respectively (**Figure 2H**). In the oxic DCM layer, metabolite concentrations were 1.8-fold higher at the offshore ODZ station and 4.8-fold higher at the coastal ODZ station compared to the NPG. Within each ODZ profile, metabolite carbon concentrations in the anoxic SCM were ∼3.4-3.6 times lower than in the oxic DCM. While metabolite concentrations were generally higher at ODZ sites and often had secondary maxima at the SCM, they decreased significantly with depth below the euphotic zone.

A core set of metabolites consistently dominated the particulate pools across both NPG oxic and ODZ stations, suggesting a fundamental similarity in chemical composition between environments (**Figure 2C, 2F, 2I; Figures S2, S3**). The most abundant compounds identified included osmolytes such as GBT, trimethylamine-N-oxide (TMAO), homarine, gonyol, sucrose, and trehalose; the amino acids glycine and glutamic acid; and the nucleobases thymine and guanine. GBT, an N-containing osmolyte, was the most abundant metabolite measured in each sample set, with concentrations as high as 15 nM in the coastal ODZ oxic DCM **(Figure S2, S4)**. At the ODZ offshore and coastal stations, GBT comprised up to ∼1.8% of the particulate carbon pool in the DCM, compared to only 0.5% at the oxic NPG DCM **(Figure 3)**. The osmolyte TMAO was detected at concentrations of up to ∼4 nM, while homarine and the S-containing metabolite gonyol were observed at levels up to ∼3 nM and ∼1 nM, respectively, within the oxic surface layers (**Figure S4**). Additionally, sucrose was up to ∼0.3 nM, and trehalose was ∼0.05 nM in the same layers. Among amino acids, glycine and glutamic acid were the most abundant, the latter of which can also function as an osmolyte, reaching ∼25 nM and ∼14 nM, respectively, in the coastal ODZ oxic surface (**Figure S6**). Nucleobases were distributed throughout the upper portion of the water columns, with thymine, a pyrimidine, detected at concentrations of up to ∼4.5 nM (**Figure S7**). Together, the data highlight the stable contribution of these metabolites to upper euphotic zone particulate carbon.

**Figure 3:**
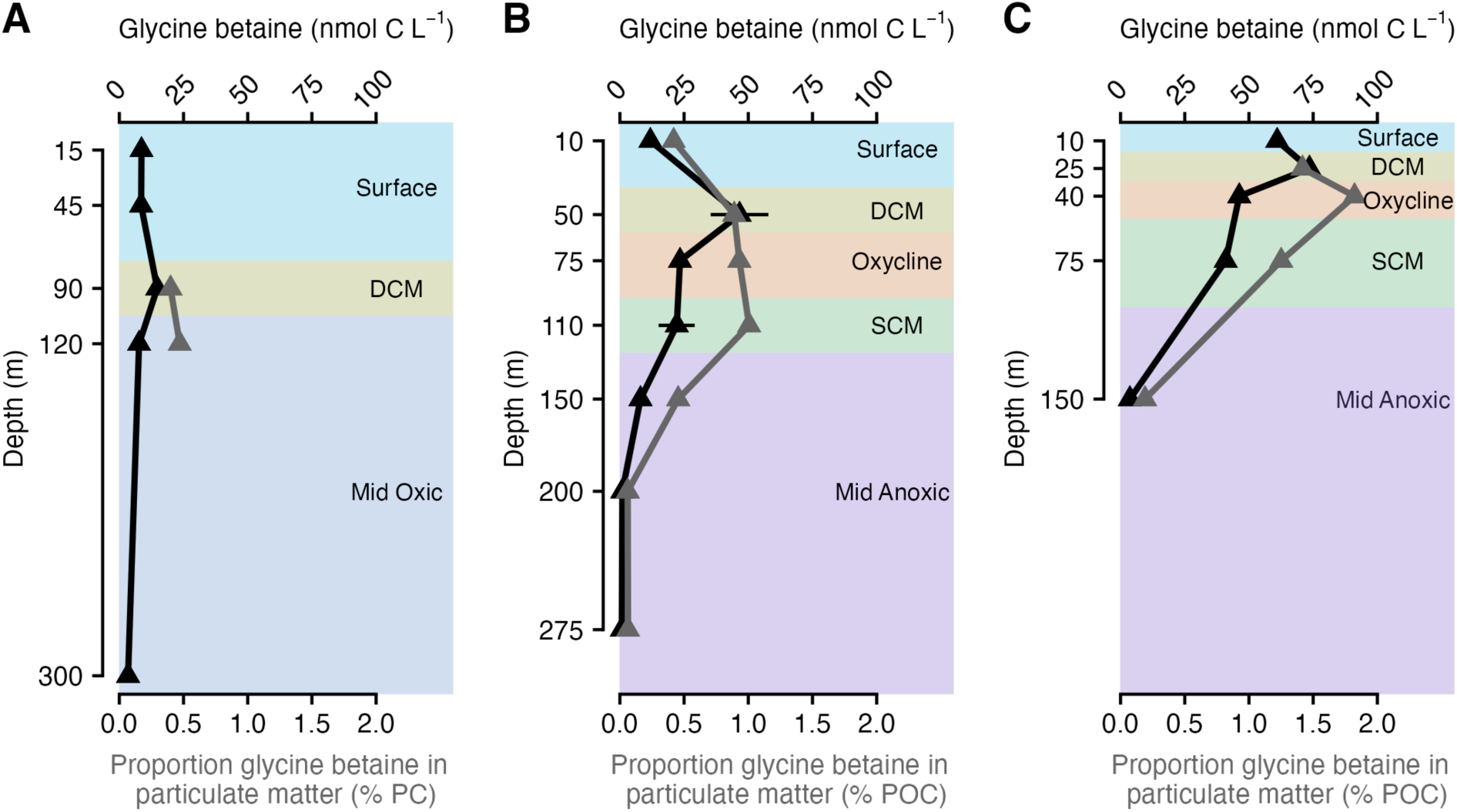
Glycine betaine depth profiles as concentration of carbon and proportion of particulate carbon (PC) particulate organic carbon (POC) from the oxic NPG **(A)**, offshore ODZ (St. P2) **(B)**, and coastal ODZ (St. P1) **(C).**

While most metabolites peaked in the oxic surface or DCM, several also exhibited secondary maxima at the SCM or in deeper mesopelagic layers (**Figures S4–S7**). At the ODZ stations, the osmolytes GBT, sucrose, and trehalose reached maximum concentrations in the surface or DCM but also had secondary peaks at the SCM or below (**Figure S4, S5**). Within the ODZ SCM, GBT made up ∼1.2% of the particulate carbon pool at the offshore and coastal stations **(Figure 3)**. Sucrose persisted at ∼0.05-0.19 nM in the SCM at the ODZ stations, making up nearly 0.4% of the POC pool at the coastal ODZ SCM. TMAO was below detection within the ODZ at both stations but was measurable (as high as ∼0.25 nM) below the ODZ in the deep oxycline (1000m and 1500m) at the coastal ODZ station (**Figure S8**). The osmolyte arsenobetaine was among the few metabolites measured with maxima below the surface and DCM layers, with concentrations over ∼0.004 nM at the SCM of both ODZ stations and at 300 m in the oxic NPG station (**Figure S5**). Additionally, amino acids such as glutamic acid and aspartic acid, had minor peaks within the SCM as well **(Figure S6)**. Nucleobase profiles followed profiles of particulate organic carbon concentrations closely, peaking at the DCM at the NPG station and in the oxic surface or DCM layers, in addition to the SCM, at the ODZ stations. Complementary base-pairs (adenine with thymine, and guanine with cytosine) showed similar depth-related trends (**Figure S7**). These patterns suggest that while most metabolites are concentrated in the upper euphotic zone, select compounds, including arsenobetaine and TMAO, persist or peak in deeper mesopelagic waters.

Multivariate patterns in metabolite composition were influenced more strongly by depth than by station. Non-metric multidimensional scaling (NMDS) was used to visualize intrinsic patterns in the quantified metabolite pools, assigning equal weight to each metabolite. The resulting ordination revealed a shift in metabolite composition across the seven environmentally defined depth groupings at each station (**Figure S9**, ANOSIM stat = 0.681, *p* = 0.001). The NMDS ordination also suggested some differences between stations (**Figure S9**, ANOSIM stat = 0.086, *p* = 0.039), but the low ANOSIM statistic implies that dissimilarities are greater between the groups than between the stations. The oxic surface and DCM samples clustered separately between stations, which is consistent with the high variability in total metabolite abundance between sample sites. The oxycline and SCM comprised a tight range of multivariate space, further indicating that these waters are very similar despite being sampled from both coastal and offshore ODZ stations. The mid-oxic samples of the NPG station are within the same multivariate space as the mid-anoxic samples of the ODZ stations, suggesting that the measured metabolite pools are similar in the mesopelagic. A dendrogram based on Euclidian distance was also used to visualize clustering of both samples and metabolites (**Figure S10**).

Indicator species analysis was used to identify metabolites associated with each of the seven environmentally defined depth groups (**Table S4**). Most quantified metabolites were strongly and significantly associated with a combination of the oxic surface and DCM. These include the N-containing amino acids glutamine, aspartic acid, and arginine, S-containing osmolytes DMSP and gonyol, and the sugar sucrose (**Table S4**). However, only a handful of metabolites acted as indicators across groups defined by anoxic waters. Specifically, six compounds (GBT, creatine, ectoine, tryptophan, and the nucleobases thymine and cytosine) were indicative of the SCM. Several compounds, including arsenobetaine, glutamic acid, key intermediates of the citric acid cycle, and the nucleobases adenine, thymine, and cytosine, were associated with mid-anoxic depths (**Table S4**). Overall, most indicator metabolites were found in the upper oxic layers, with only select compounds associated with deeper anoxic waters.

### Glycine betaine synthesis and transformation pathways

Given that GBT was the most abundant metabolite measured and was an indicator of the SCM, we investigated which organisms might be responsible for the active production and transformation of GBT along the oxycline, SCM, and mid-anoxic depths by searching metatranscriptomes collected from the offshore and coastal ODZ sample sites for genes encoding GBT related proteins in KEGG (**Table S6**). This revealed three pathways of GBT metabolism expressed in both ODZ profiles, with the overall highest expression levels at the coastal ODZ station. We found two active GBT production pathways: (1) the choline oxidation pathway from choline and betaine aldehyde and (2) the methylation pathway from glycine sarcosine and dimethylglycine (**Figure 4**). We also found expression of a GBT demethylation pathway into dimethyl glycine, sarcosine, and glycine. We explored other known pathways of GBT transformation, including the GBT reduction pathway into trimethylamine (TMA) and TMAO (by *grdH*) and the choline lipid pathway into choline (by *ATRR*). However, these pathways were not represented in the metagenomes (**Table S6**), and thus were likely not active pathways for GBT production and transformation in the oxycline and ODZ. This analysis enabled taxonomic resolution of GBT production and transformation pathways in the ODZ.

**Figure 4:**
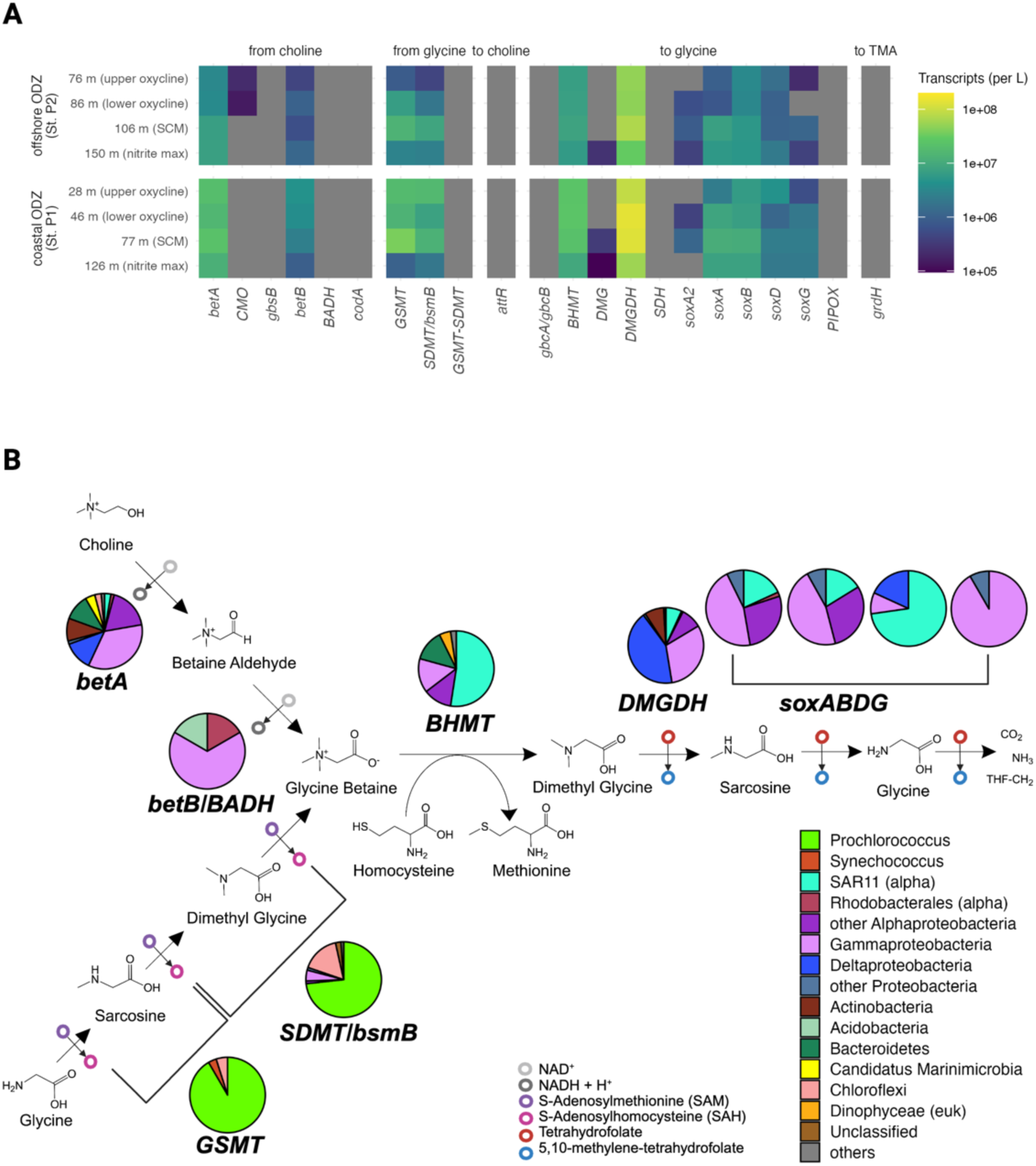
Glycine betaine synthesis and transformation expression patterns at the ODZ stations. **A)** Abundance of transcripts with depth at the offshore (P2) and coastal (P1) ODZ stations. Genes with expression include *betA*, choline dehydrogenase; *CMO*, choline monooxygenase; *betB*, betaine aldehyde dehydrogenase; *GSMT*, glycine-sarcosine methyltransferase; *bsmB/SDMT*, sarcosine dimethyltransferase; *BHMT*, betaine-homocysteine methyltransferase; DMG, dimethylglycine oxidase; *DMGDH*, dimethylglycine dehydrogenase; *SoxABDG*, heterotetrameric sarcosine oxidase. Genes not present in metagenomes and without expression include *gbsB*, choline dehydrogenase; *BADH*, betaine aldehyde dehydrogenase*; codA*, choline oxidase; *GSMT-SDMT*, glycine/sarcosine/dimethylglycine N-methyltransferase; *attR*, glycine betaine reductase; *gbcA/gbcB*, glycine betaine monooxygenase; *SDH*, sarcosine dehydrogenase; *PIPOX*, sarcosine oxidase; *grdH*, betaine reductase. Pathway details and KEGG numbers for each gene provided in Table S6. **B)** Glycine betaine (GBT) metabolic pathways expressed in the secondary chlorophyll maximum (SCM) at the offshore ODZ station (P2) with proportional taxonomy for each gene shown. Note pies are of equal size for each gene even though overall transcript abundance for each gene differed as in A.

The choline oxidation pathway for GBT synthesis was expressed in the oxycline, SCM, and mid-anoxic depths at both the offshore and coastal ODZ sample sites (**Figure 4A**). At the SCM of the offshore ODZ station, the choline oxidation pathway from choline to betaine aldehyde, catalyzed by an NAD^+^-dependent choline dehydrogenase (*betA*), and was primarily expressed by Alphaproteobacteria, and Gammaproteobacteria (**Figure 4B**). The second reaction step, catalyzed by an NAD^+^-dependent betaine aldehyde dehydrogenase (*betB*), was dominated by Gammaproteobacteria. At both stations the highest expression for these genes was in the SCM (**Figure 4A, S11)** and taxonomic contributions to these pathways underwent shifts along the oxygen gradients. For instance, Alphaproteobacteria and Gammaproteobacteria contribution to *betA* expression increased at anoxic depths, where Deltaproteobacteria contribution to *betA* expression decreased. Chloroflexi expressed *betA* in the oxycline at both ODZ stations. Expression of *betB* shifted from primarily Gammaproteobacteria and Actinobacteria in the upper oxycline to Gammaproteobacteria and Acidobacteria at the SCM and below (**Figure S11**). While Gammaproteobacteria were the main contributors to both *betA* and *betB* expression, the relative contribution of other taxa shifted with depth and oxygen availability.

The expression of the methylation pathway for GBT synthesis was also highest in the SCM (**Figure 4A**) and almost entirely attributed to *Prochlorococcus*, with smaller contributions from *Synechococcus* and Chloroflexi (**Figure 4B**, **S11**). In this pathway, GBT is synthesized from glycine by three methylation reactions with S-adenosylmethionine as methyl donor, and sarcosine and dimethylglycine serving as the intermediates. Glycine sarcosine methyltransferase (GSMT) catalyzes the methylation of glycine or sarcosine to form sarcosine or dimethylglycine, and sarcosine dimethylglycine methyltransferase (*SDMT*/*bsmB*) catalyzes the methylation of dimethylglycine to GBT, respectively. The coastal ODZ station had the highest expression levels overall, however there were slight differences in expression patterns across the oxygen gradients at each ODZ station. The coastal ODZ station had higher expression for *GSMT* and *SDMT/bsmB* within the upper oxycline and lower expression within the mid-anoxic depth compared to the offshore ODZ station. At the offshore ODZ Station, *Prochlorococcus* express *GSMT* and *SDMT/bsmB* as deep as 150 meters depth at the bottom of the SCM. These patterns highlight *Prochlorococcus* as the primary contributor to GBT synthesis via methylation in the SCM and upper oxycline.

GBT synthesis via the methylation pathway at the coastal and offshore ODZ stations was dominated by transcripts from *Prochlorococcus* ecotypes specific to the ODZ (**Figure 5**). Screening of publicly available marine picocyanobacterial genomes revealed that only a subset contained the genetic potential for GBT production. A maximum likelihood phylogenetic tree of *SDMT/bsmB* from these genomes (Figure 5A) supports the interpretation that this trait is vertically inherited within these lineages, rather than acquired via horizontal gene transfer. Within the *Prochlorococcus* group, the genetic potential for GBT synthesis is restricted to the LL IV, AMZ Ia and Ib (LL V), AMZ II (LL VI) and AMZ III clades. Along the oxycline, SCM, and mid-anoxic depths of the offshore and coastal ODZ stations, these *Prochlorococcus* lineages dominate the expression of *SDMT/bsmB* (**Figure 5B, 5C)**. However, there were variations in expression among clades along the oxygen gradients observed at both the coastal and offshore ODZ stations. Specifically, the lower oxycline and anoxic waters had *SDMT/bsmB* expression by primarily AMZ Ia and AMZ Ib *Prochlorococcus*. In the upper oxycline of the coastal ODZ station, which overlaps with the DCM, *SDMT/bsmB* expression was dominated by AMZ II *Prochlorococcus* (**Figure 5C)**. Thus, *SDMT/bsmB* expression may reflect ecotype-specific activity along oxygen gradients.

**Figure 5:**
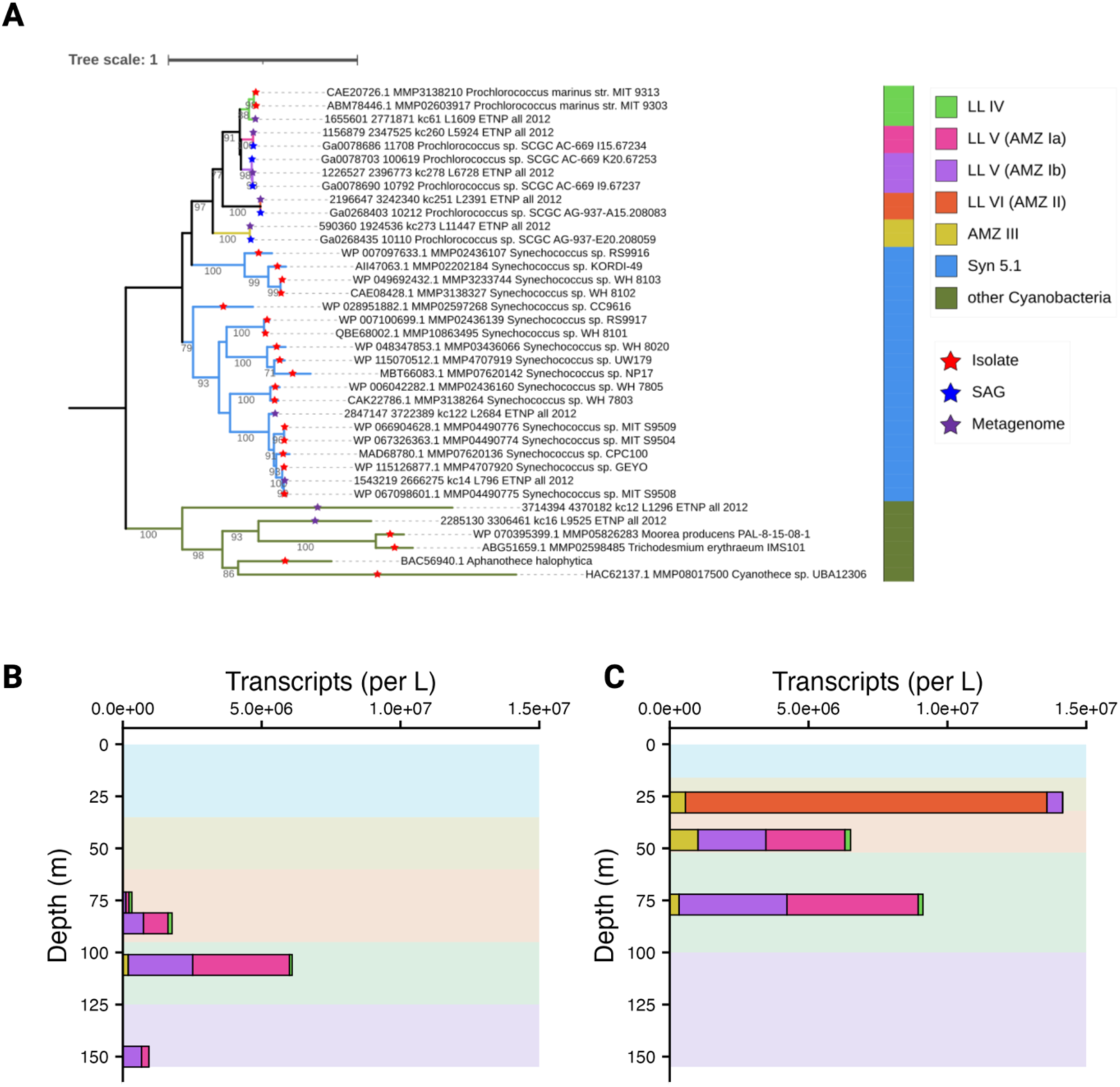
*Prochlorococcus* diversity and expression of the glycine betaine synthesis gene *SDMT/bsmB* in the ODZ (**A**) Maximum likelihood phylogenetic tree of *SDMT/bsmB* in representative Cyanobacteria. Names with ETNP indicate sequences assembled from 2012 metagenome at offshore station (P2). Bootstrap values ≥60 are shown. Colored shading indicates the taxonomic group. Expression of *SDMT/bsmB* by *Prochlorococcus* with depth at the offshore (**B**) and coastal (**C**) ODZ stations.

The GBT demethylation pathway, which consumes GBT, was expressed across all depths measured at both the offshore and coastal ODZ sample sites (**Figure 4A**) and involved a consortium of different organisms (**Figure 4B, S11).** At the SCM of the offshore station betaine-homocysteine methyltransferase (*BHMT*) expression, the first step of GBT demethylation that is predicted to convert GBT and homocysteine to dimethylglycine and methionine in marine bacteria, was dominated by SAR11 and other Alphaproteobacteria, with contribution from Gammaproteobacteria, Bacteroidetes, and dinoflagellates (**Figure 4B**). The subsequent demethylation of dimethyl glycine to sarcosine through dimethylglycine oxidase (*DMGDH*) had the most abundant transcript among these pathways and was dominated by Deltaproteobacteria and Gammaproteobacteria. The third step of the demethylation pathway, catalyzed by sarcosine dehydrogenases (*soxBDAG* or *SDH*), was expressed by a variety of Proteobacteria. SAR11 expressed the entire GBT transformation pathway (all three steps) at both ODZ stations (**Figure S12**).The expression patterns of specific taxa in these pathways with depth were consistent across both ODZ stations, with higher expression levels at the coastal ODZ station (**Figure S11,**). At both ODZ stations, SAR11 consistently expressed all steps of the GBT demethylation pathway across sampled depths (**Figure S12**).

## Discussion

This study presents depth-resolved profiles of particulate metabolites in an oxygen-deficient zone (ODZ), providing a new view of microbial physiology and chemical interactions in low-oxygen marine environments. By linking metabolite distributions to gene expression across the oxycline, secondary chlorophyll maximum (SCM), and anoxic layers, we reveal taxon-specific pathways for glycine betaine (GBT) production and consumption. These findings offer a detailed, community-level perspective on how GBT connects phototrophic and heterotrophic metabolisms across oxygen gradients. In contrast, other osmolytes such as trimethylamine-N-oxide (TMAO) were detected only in oxygenated waters and were absent from the anoxic core, reflecting differences in compound distributions across redox boundaries. Together, these results expand our understanding of the metabolic landscape in ODZs and identify GBT as a central metabolite in microbial interactions.

Intracellular metabolites provide a window into the metabolic activity and physiological state of microbial communities, offering context for interpreting patterns in carbon cycling and productivity. In this study, the 87 measured particulate metabolites accounted for up to 5.5% of particulate carbon, with the highest total concentrations of metabolites typically observed in the oxic surface or DCM layers (**Figure 2B, 2E, 2H**; **Table S3**). These findings align with previous observation in the North Pacific Ocean, where measured particulate metabolites comprised approximately 3-10% of particulate carbon pools in surface waters (Boysen et al., 2021; Heal et al., 2021). The elevated concentrations of surface particulate metabolites and particulate organic carbon at the ODZ stations may indicate higher primary productivity compared to the NPG station, which is situated at the northern edge of an oligotrophic gyre (**Figure 1**). Indeed, the strongest upwelling wind conditions typically occur along the west coast of Mexico in April-June (Bulgakov et al., 2005; González-Rodríguez et al., 2012), when we sampled in the ODZ, and satellite chlorophyll concentrations were relatively high at the coastal station (**Figure 1**). While both the coastal and offshore ODZ stations show comparable particulate organic concentrations in the DCM (∼5 µM), the coastal station has double the particulate metabolite concentrations and higher satellite chlorophyll concentrations in the surface (**Figure 1**) and chlorophyll fluorescence in the mixed layer (**Figure 2D, 2G**). This higher contribution of metabolites to particulate organic carbon at the coastal station most likely stem from a higher proportion of biomass relative to detritus at this location but could also be influenced by cells accumulating higher concentrations of intracellular metabolites, possibly facilitated by spring bloom conditions (Boysen et al., 2021).

The oxic surface and DCM layers at all stations were characterized by a conserved suite of abundant particulate metabolites, suggesting broadly similar phytoplankton communities (**Figure 2C, 2F, 2I**). Most metabolites were strongly and significantly associated with a combination of the oxic surface and DCM layers (**Table S4**). These included compounds typically associated with eukaryotic phytoplankton such as the S-containing osmolytes DMSP and gonyol as well as the N-containing osmolyte homarine. DMSP production and accumulation have been documented in dinoflagellates, coccolithophores, and diatoms (Keller et al., 1989; Kettles et al., 2014), as well as in certain bacterial species (Zheng et al., 2020). Gonyol is known to accumulate in culture strains of dinoflagellates and haptophytes (Nakamura et al., 1993; Heal et al., 2021). Homarine constitutes up to 0.2% of the particulate carbon of *Synechococcus* culture strains (Heal et al., 2021) and has also been detected in diatoms and haptophytes. The oxic surface and DCM waters of both the NPG and ODZ regions are expected to contain a diverse community of phytoplankton, including photosynthetic haptophytes, cryptophytes, dinoflagellates, diatoms, and picocyanobacteria (Lavin et al., 2010; Parris et al., 2014; Dutkiewicz et al., 2020; Fuchsman et al., 2022; Xu et al., 2022). The observed trends in sulfur compounds and N-containing homarine (**Figure S4**) suggest that the presence of eukaryotic phytoplankton and *Synechococcus* leaves a distinct chemical signature in the oxic surface and DCM layers.

In deeper waters, overall metabolite diversity was reduced, but select compounds such as trimethylamine-N-oxide (TMAO) displayed consistent and oxygen-sensitive depth distributions. The mid-oxic samples of the NPG station and the mid-anoxic and deep oxic samples of the ODZ stations all clustered together in the 2-dimensional NMDS plot (**Figure S9**), suggesting that the measured metabolite pools are similar within microbial communities at deeper depths. However, the osmolyte TMAO emerged as a consistent indicator of all oxic depths across stations, including the deep oxic zone below the ODZ. Interestingly, TMAO was below detection within the anoxic layers of the ODZs, reappearing within the deep oxycline (**Figure S8**). TMAO is a major osmolyte found in a variety of marine organisms and can be synthesized by marine Roseobacter and SAR11 (Chen et al., 2011; Chen, 2012; Li et al., 2017; Gao et al., 2022). TMAO can be metabolized to small, methylated amines or used as a terminal electron acceptor during anaerobic microbial respiration (Halsey et al., 2012; Lidbury et al., 2014), or used as a nitrogen source for growth (Gao et al., 2022). Under aerobic conditions, TMAO is known to be produced from choline and GBT (Sun et al., 2019). It is possible that lack of production or efficient use of TMAO as a C and N and/or energy source by SAR11 and other heterotrophs is responsible for undetectable TMAO concentrations within oxygen-deficient waters. TMAO production is not exclusive to bacteria but has also been associated with marine animals such as copepods (Strom, 1979) and various fish species (Maker et al., 1963; Goldstein and Funkhouser, 1972; Seibel and Walsh, 2002; Treberg and Driedzic, 2002; Bockus and Seibel, 2016). In these animals, TMA produced by gut microbiota or obtained from the environment is detoxified into TMAO. The oxygen-deficient layers in ODZs may disrupt the vertical migration (Maas et al., 2012, 2014; Wishner et al., 2018) or reduce the metabolic activity (Kiko et al., 2016) of animals. This could limit the release of TMAO by such animals, potentially contributing to the pronounced disappearance of TMAO in ODZs as well.

The secondary chlorophyll maximum (SCM) is a distinct feature within the ODZ, characterized by anoxia, low light, high nutrients, and a microbial community in which *Prochlorococcus* dominates the photosynthetic population. The SCM samples from the coastal and offshore ODZ clustered together in the 2-dimensional NMDS plot (**Figure S9**), demonstrating reproducibility of SCM particulate metabolite pools within the ODZ system (**Table S1**). Compared to the DCM, the SCM contains a much less diverse phytoplankton community, one almost entirely dominated by the picocyanobacteria *Prochlorococcus* rather than photosynthetic eukaryotes (Johnson et al., 1999; Goericke et al., 2000; Aldunate et al., 2022; Fuchsman et al., 2022; Wong et al., 2023). Thus, the SCM provides an opportunity to observe the chemical signature of ODZ ecotypes of *Prochlorococcus.* The most abundant metabolites in the SCM included GBT, sucrose, trehalose, nucleobases, as well as the amino acids aspartic acid and glutamic acid. Glutamic acid, among other amino acids, serves as a common compatible solute in a wide variety of marine microorganisms (Measures, 1975; Csonka, 1989; Welsh, 2000; Brill et al., 2011; Durham et al., 2019) and aligns with general biomass trends (Johnson et al., 2023). However, GBT and sucrose along with glutamic acid and aspartic acid, are known to accumulate in *Prochlorococcus* cells as osmolytes (Klähn et al., 2010; Durham et al., 2019; Boysen et al., 2021; Heal et al., 2021; Kujawinski et al., 2023), potentially contributing to the observed pools within the SCM. Of these abundant compounds, only GBT was an indicator of the SCM. Among measured metabolites, the SCM had the highest proportion of GBT across all depths and stations (**Figure 2**), contributing to ∼1.2% of POC (**Figure 3**). These findings identify GBT as a defining metabolite in SCM waters dominated by ODZ ecotypes of *Prochlorococcus*.

The glycine methylation pathway to produce GBT, encoded by *GSMT* and *SDMT/bsmB*, is predominately expressed by *Prochlorococcus* in the SCM (**Figure 4**). This pathway has been characterized in several halophilic phototrophic bacteria and methanogenic archaea (Lai et al., 1999; Nyyssölä et al., 2000; Waditee et al., 2003; Yang et al., 2022) as well as in *Synechococcus* (Lu et al., 2006). In *Synechococcus* and *Prochlorococcus* LL IV isolates the *GSMT* and *SDMT/bsmB* genes are found co-located on the genome along with three genes that encode an ABC transporter for GBT (*proVWX*), which may be used for reuptake of GBT that leaks into the periplasmic space (Scanlan et al., 2009). Both *Synechococcus* and *Prochlorococcus* LL IV isolates have been observed to accumulate GBT (Klähn et al., 2010; Kujawinski et al., 2023). Among cultured *Prochlorococcus* isolates, GBT production appears to be restricted to the LL IV clade (Scanlan et al., 2009; Klähn and Hagemann, 2011; Kujawinski et al., 2023). Here, we show that the uncultured ecotypes within the ODZ (AMZ Ia, AMZ Ib, AMZ II, and AMZ III) all also have the genetic potential for GBT production (**Figure 5A**). Like LL IV *Prochlorococcus* and *Synechococcus*, they possess a genomic cluster of GBT related genes, including both *GSMT* and *SDMT/bsmB* and a GBT ABC transporter (*proVWX*). The AMZ clades are more deeply branching within the *Prochlorococcus* group, and this phylogenetic pattern suggests that the GBT synthesis genes were not acquired recently through horizontal gene transfer, but instead retained from a common ancestor shared with *Synechococcus*. These *Prochlorococcus* ecotypes may accumulate GBT, a quaternary ammonium compound, as an adaptation to the nitrogen-rich, anoxic environment in which they thrive. With the exception of the use of amino acids as osmolytes, other *Prochlorococcus* ecotypes use nitrogen-free osmolytes such as sucrose and glucosylglycerol (Hagemann, 2011; Heal et al., 2021). The loss of the GBT synthesis pathway in other *Prochlorococcus* ecotypes coincides with genome reduction during adaptation to nitrogen-poor waters higher in the water column (Rocap et al., 2003; Kettler et al., 2007; Berube et al., 2015; Braakman et al., 2017).

ODZ *Prochlorococcus* contribute directly to the accumulation of GBT within the oxycline and SCM, with distinct ecotypes dominating GBT synthesis in different parts of the water column (**Figure 5B**). The most abundant *Prochlorococcus* ecotypes within the ODZ, AMZ Ia and AMZ Ib, (Lavin et al., 2010; Fuchsman et al., 2021; Ulloa et al., 2021; Jaffe et al., 2024), exhibit the highest *SDMT/bsmb* activity in anoxic waters. In contrast, in the upper oxycline of ODZ coastal station P1, AMZ II *Prochlorococcus* dominates *SDMT/bsmb* expression. GBT degradation or transformation genes are not found in *Prochlorococcus*, suggesting that these ecotypes produce GBT to function as a compatible solute and likely accumulate high amounts of GBT within the oxycline and SCM, contributing to the GBT pools there (**Figure 3**). Once synthesized, *Prochlorococcus* appear unable to further metabolize GBT, making GBT available for release into the environment through excretion or via viral-induced cell lysis (Fuchsman et al., 2019, 2021).

Other contributions to the GBT pools across the oxycline and SCM were made via the choline oxidation pathway. This transformation begins with the already fully methylated substrate, choline, which is oxidized to betaine aldehyde and then to GBT. This pathway was primarily expressed by Alphaproteobacteria and Gammaproteobacteria (**Figure 4**). Evidence from comparative genomics suggests that over half of cultivated marine Alphaproteobacteria and Gammaproteobacteria strains have the genomic potential for GBT synthesis through choline oxidation (McParland et al., 2021). Cultured SAR11 strains have the genetic potential for the first step of the oxidation pathway but are missing the last step where betaine aldehyde is converted to GBT (Haro-Moreno et al., 2019). Gammaproteobacteria had the highest expression of *betB* in our ODZ samples overall. Thus, in the anoxic core of the ODZ, well below depths where *Prochlorococcus* are found, the choline oxidation pathway is likely the primary source of GBT, and Gammaproteobacteria may be the dominant producer.

To explore GBT fate and potential sinks in the ODZ, we examined expression of demethylation pathways (Boysen et al., 2022). Using our search criteria, the expression of *BHMT*, the first step of GBT demethylation that is predicted to convert GBT and homocysteine to dimethylglycine and methionine in marine bacteria, was dominated by SAR11 and possibly other Alphaproteobacteria, with contributions from Gammaproteobacteria, Bacteroidetes, and dinoflagellates (**Figure 4**). SAR11 expressed transcripts for all three known demethylation steps (**Figure S12**), though a few other taxa may also possess partial or complete pathways. Additionally, downstream enzymes such as *DMGDH* were expressed in organisms lacking detectable *BHMT*, suggesting that alternative or uncharacterized demethylation pathways may exist. These patterns also raise the possibility that GBT-derived intermediates are released or shared among different members of the community, allowing multiple organisms to participate in the stepwise transformation of GBT. Together, these findings highlight the complexity of GBT cycling, with production and degradation partitioned across taxa and depths, and its breakdown potentially involving collaborative processing within the microbial community.

GBT is a highly available substrate that can meet many of the metabolic needs of SAR11. SAR11 was first cultivated in minimal media containing methionine, pyruvate, and glycine, suggesting a need for these substrates (Carini et al., 2012). SAR11 cells possess a high affinity and multifunctional GBT transporter (Noell and Giovannoni, 2019; Mausz et al., 2022), suggesting that SAR11 can readily take up GBT in the environment. Once inside the cell, GBT not only meets SAR11’s specific glycine requirement (Tripp, 2013), as shown in LL IV *Prochlorococcus* co-culture studies (Becker et al., 2019), but also serves as a key precursor for the essential amino acid methionine. SAR11 cells lack the cobalamin (Vitamin B_12_)-dependent (*metH*) and cobalamin-independent (*metE*) pathways for methionine synthesis, which is thought to make them reliant on external methionine sources (Giovannoni, 2017). However, SAR11 can use GBT to produce methionine through the putative *BHMT* pathway, where a GBT methyl group is transferred to homocysteine, resulting in methionine (Sun et al., 2011) (**Figure 4B**). This capability may allow SAR11 to bypass the need for *metH* and *metE,* with *BHMT* functioning as a methionine synthase. Although in LL IV *Prochlorococcus* co-culture studies SAR11 was unable to grow in the absence of methionine (Becker et al., 2019), this may have been due to the lack of an alternate source of reduced sulfur in the minimal medium. Methylated substrates like GBT reduce the lag phase of starved heterotrophic bacteria (Narváez-Barragán et al., 2024), suggesting that GBT may support the growth and survival of SAR11 during periods of fluctuating resource availability. Additionally, the remaining methyl groups from GBT are predicted to be transferred to tetrahydrofolate in reactions that are coupled to partial oxidation and further conversion to CO_2_, potentially acting as an energy source (Sun et al., 2011; Lidbury et al., 2015). SAR11 is strongly associated with GBT uptake in both coastal (Noell and Giovannoni, 2019; Mausz et al., 2022) and open ocean (Boysen et al., 2022) waters. The widespread presence of GBT in marine environments and the ability of SAR11 to synthesize methionine and glycine from GBT, including in ODZs, is likely crucial to the ecological success of SAR11.

## Conclusion

In this study, we present particulate metabolite profiles in an ODZ. Depth profiles of the 87 measured metabolites were largely congruent with those observed in the oxic oligotrophic ocean, with a few key differences: the absence of TMAO in anoxic waters and the enrichment of GBT at the SCM. The unique phototrophic community composition of the SCM provides an opportunity to observe the chemical signature of ODZ ecotypes of *Prochlorococcus*, which contribute a substantial portion of in situ organic carbon within and below the SCM (Fuchsman et al., 2019; Aldunate et al., 2022). The release of GBT by *Prochlorococcus* cells, whether through excretion or viral-induced cell lysis (Fuchsman et al., 2019, 2021), likely serves as a significant source of GBT to a wide range of co-occurring heterotrophs, including SAR11. With its streamlined genome, SAR11 lacks traditional methionine synthesis pathways, and may exploit GBT for methionine production via the putative *BHMT* pathway. The acquisition of GBT for methionine production by SAR11 may also occur deeper in the anoxic ODZ, where it is produced by Gammaproteobacteria, as well as in oxic waters worldwide, where it is produced by a consortium of other heterotrophs and phototrophs. This underscores the critical role of GBT as a metabolic currency mediating microbial interactions in the marine environment and supporting the growth of SAR11, the most numerically abundant heterotroph on earth.

## Acknowledgements

We thank the captains and crew of the R/V *Revelle* and R/V *Marcus Langseth* for their invaluable support during our research expeditions. Special thanks to Angela Boysen and Katherine Heal for their assistance with sample collection, metabolomics pipeline development and data analysis. We are grateful to Angelique White for providing particulate carbon data from the North Pacific Gyre to Cedar McKay for metagenomic assembly, Timothy Mattes and Susan Burke for collecting transcript samples, Timothy Mattes for sequencing the transcripts, and Zeta Lai for her contributions to researching metabolic pathways.

Support for this work was provided by the National Science Foundation (DEB-1542240 and OCE-2022911 to G.R., Award ID 2125886 to A.E.I., OCE-2201310 to R.M.M., and a Graduate Research Fellowship to N.A.K.), the Simons Foundation (SCOPE Award ID 329108 and Award ID 385428 to A.E.I.), and Horn Point Laboratory start-up funds to C.A.F.

Conflict of Interest: none declared

## Data Availability Statement

CTD and nutrient data are available at BCO-DMO dataset 779185.

Transcripts can be downloaded from BioProject PRJNA727903 or MG-RAST project mgp92168.

Metagenomes can be downloaded from Bioproject PRJNA350692.

Metabolomic data is available at Metabolomics Workbench PR002263

(https://www.metabolomicsworkbench.org/)

## Citations

Aldunate, M., P. von Dassow, C. A. Vargas, and O. Ulloa. 2022. Carbon Assimilation by the Picoplanktonic Community Inhabiting the Secondary Chlorophyll Maximum of the Anoxic Marine Zones of the Eastern Tropical North and South Pacific. Frontiers in Marine Science 9.

Altschul, S. F., T. L. Madden, A. A. Schäffer, J. Zhang, Z. Zhang, W. Miller, and D. J. Lipman. 1997. Gapped BLAST and PSI-BLAST: a new generation of protein database search programs. Nucleic Acids Res. 25: 3389–3402.

Aramaki, T., R. Blanc-Mathieu, H. Endo, K. Ohkubo, M. Kanehisa, S. Goto, and H. Ogata. 2020. KofamKOALA: KEGG Ortholog assignment based on profile HMM and adaptive score threshold. Bioinformatics 36: 2251–2252.

Azam, F., T. Fenchel, J. Field, J. S. Gray, L. Meyer, and T. F. Thingstad. 1983. The Ecological Role of Water-Column Microbes in the Sea. Marine Ecology Progress Series 10: 257–263.

Banse, K., S. W. A. Naqvi, and J. R. Postel. 2017. A zona incognita surrounds the secondary nitrite maximum in open-ocean oxygen minimum zones. Deep Sea Research Part I: Oceanographic Research Papers 127: 111–113.

Becker, J. W., S. L. Hogle, K. Rosendo, and S. W. Chisholm. 2019. Co-culture and biogeography of Prochlorococcus and SAR11. The ISME Journal 13: 1506–1519.

Beman, J. M., and M. T. Carolan. 2013. Deoxygenation alters bacterial diversity and community composition in the ocean’s largest oxygen minimum zone. Nature Communications 4: 2705.

Berube, P. M., S. J. Biller, A. G. Kent, J. W. Berta-Thompson, S. E. Roggensack, K. H. Roache-Johnson, M. Ackerman, et al. 2015. Physiology and evolution of nitrate acquisition in Prochlorococcus. The ISME Journal 9: 1195–1207.

Bligh, E. G., and W. J. Dyer. 1959. A RAPID METHOD OF TOTAL LIPID EXTRACTION AND PURIFICATION. Canadian Journal of Biochemistry and Physiology 37: 911–917.

Bockus, A. B., and B. A. Seibel. 2016. Trimethylamine oxide accumulation as a function of depth in Hawaiian mid-water fishes. Deep Sea Research Part I: Oceanographic Research Papers 112: 37–44.

Boyer, T. P., H. E. García, R. A. Locarnini, M. M. Zweng, A. V. Mishonov, J. R. Reagan, K. A. Weathers, et al. 2018. World Ocean Atlas 2018. [oxygen].

Boysen, A. K., L. T. Carlson, B. P. Durham, R. D. Groussman, F. O. Aylward, F. Ribalet, K. R. Heal, et al. 2021. Particulate Metabolites and Transcripts Reflect Diel Oscillations of Microbial Activity in the Surface Ocean. mSystems 6: 10.1128/msystems.00896-20.

Boysen, A. K., B. P. Durham, W. Kumler, R. S. Key, K. R. Heal, L. T. Carlson, R. D. Groussman, et al. 2022. Glycine betaine uptake and metabolism in marine microbial communities. Environmental Microbiology 24: 2380–2403.

Boysen, A. K., K. R. Heal, L. T. Carlson, and A. E. Ingalls. 2018. Best-Matched Internal Standard Normalization in Liquid Chromatography–Mass Spectrometry Metabolomics Applied to Environmental Samples. Analytical Chemistry 90: 1363–1369.

Braakman, R., M. J. Follows, and S. W. Chisholm. 2017. Metabolic evolution and the self-organization of ecosystems. Proceedings of the National Academy of Sciences 114: E3091–E3100.

Brill, J., T. Hoffmann, M. Bleisteiner, and E. Bremer. 2011. Osmotically Controlled Synthesis of the Compatible Solute Proline Is Critical for Cellular Defense of Bacillus subtilis against High Osmolarity. Journal of Bacteriology 193: 5335.

Brister, J. R., D. Ako-Adjei, Y. Bao, and O. Blinkova. 2015. NCBI viral genomes resource. Nucleic Acids Research 43: D571–577.

Bulgakov, S. N., N. P. Bulgakov, E. N. Mikhailova, and N. B. Shapiro. 2005. Generation of Upwelling near the Pacific Coast of Mexico. Physical Oceanography 15: 27–36.

Carini, P., L. Steindler, S. Beszteri, and S. J. Giovannoni. 2012. Nutrient requirements for growth of the extreme oligotroph ‘Candidatus Pelagibacter ubique’ HTCC1062 on a defined medium. The ISME Journal 7: 592.

Cepeda-Morales, J., E. Beier, G. Gaxiola-Castro, M. F. Lavín, and V. Godínez. 2009. Effect of the oxygen minimum zone on the second chlorophyll maximum in the Eastern Tropical Pacific off Mexico. Ciencias Marinas 35: 389–403.

Chen, Y. 2012. Comparative genomics of methylated amine utilization by marine Roseobacter clade bacteria and development of functional gene markers (tmm, gmaS). Environmental Microbiology 14: 2308–2322.

Chen, Y., N. A. Patel, A. Crombie, J. H. Scrivens, and J. C. Murrell. 2011. Bacterial flavin-containing monooxygenase is trimethylamine monooxygenase. Proceedings of the National Academy of Sciences of the United States of America 108: 17791–17796.

Csonka, L. N. 1989. Physiological and genetic responses of bacteria to osmotic stress. Microbiological Reviews 53: 121–147.

DeVries, T., C. Deutsch, P. Rafter, and F. Primeau. 2012. Marine denitrification rates determined from a global 3-D inverse model. Biogeosciences Discussions 9: 14013–14052.

Durham, B. P., A. K. Boysen, L. T. Carlson, R. D. Groussman, K. R. Heal, K. R. Cain, R. L. Morales, et al. 2019. Sulfonate-based networks between eukaryotic phytoplankton and heterotrophic bacteria in the surface ocean. Nature Microbiology 4: 1706–1715.

Durham, B. P., A. K. Boysen, K. R. Heal, L. T. Carlson, R. Boccamazzo, C. R. Deodato, W. Qin, et al. 2022. Chemotaxonomic patterns in intracellular metabolites of marine microbial plankton. Frontiers in Marine Science 9.

Dutkiewicz, S., P. Cermeno, O. Jahn, M. J. Follows, A. E. Hickman, D. A. A. Taniguchi, and B. A. Ward. 2020. Dimensions of marine phytoplankton diversity. Biogeosciences 17: 609–634.

Edgar, R. C. 2004. MUSCLE: multiple sequence alignment with high accuracy and high throughput. Nucleic Acids Research 32: 1792–1797.

Falkowski, P. G. 1994. The role of phytoplankton photosynthesis in global biogeochemical cycles. Photosynthesis Research 39: 235–258.

Fuchsman, C. A., M. C. G. Carlson, D. Garcia Prieto, M. D. Hays, and G. Rocap. 2021. Cyanophage host-derived genes reflect contrasting selective pressures with depth in the oxic and anoxic water column of the Eastern Tropical North Pacific. Environmental Microbiology 23: 2782–2800.

Fuchsman, C. A., L. Cherubini, and M. D. Hays. 2022. An analysis of protists in Pacific oxygen deficient zones: implications for Prochlorococcus and N-producing bacteria. Environmental Microbiology 24: 1790–1804.

Fuchsman, C. A., A. H. Devol, J. K. Saunders, C. McKay, and G. Rocap. 2017. Niche Partitioning of the N Cycling Microbial Community of an Offshore Oxygen Deficient Zone. Frontiers in Microbiology 8.

Fuchsman, C. A., M. E. Duffy, J. A. Cram, P. Huanca-Valenzuela, B. P. Gregory, L. V. Plough, J. J. Pierson, et al. 2025. Contributions of Vertically Migrating Metazoans to Sinking and Suspended Particulate Matter Fuel N_2_ Production in the Eastern Tropical North Pacific Oxygen Deficient Zone. Global Biogeochemical Cycles 39: e2024GB008365.

Fuchsman, C. A., H. I. Palevsky, B. Widner, M. Duffy, M. C. G. Carlson, J. A. Neibauer, M. R. Mulholland, et al. 2019. Cyanobacteria and cyanophage contributions to carbon and nitrogen cycling in an oligotrophic oxygen-deficient zone. The ISME Journal 13: 2714–2726.

Ganesh, S., D. J. Parris, E. F. DeLong, and F. J. Stewart. 2014. Metagenomic analysis of size-fractionated picoplankton in a marine oxygen minimum zone. The ISME Journal 8: 187–211.

Gao, C., N. Zhang, X.-Y. He, N. Wang, X.-Y. Zhang, P. Wang, X.-L. Chen, et al. 2022. Characterization of the Trimethylamine N-Oxide Transporter From Pelagibacter Strain HTCC1062 Reveals Its Oligotrophic Niche Adaption. Frontiers in Microbiology 13: 838608.

Garcia-Robledo, E., C. C. Padilla, M. Aldunate, F. J. Stewart, O. Ulloa, A. Paulmier, G. Gregori, and N. P. Revsbech. 2017. Cryptic oxygen cycling in anoxic marine zones. Proceedings of the National Academy of Sciences 114: 8319–8324.

Giovannoni, S. J. 2017. SAR11 Bacteria: The Most Abundant Plankton in the Oceans. Annual Review of Marine Science 9: 231–255.

Goericke, R., R. J. Olson, and A. Shalapyonok. 2000. A novel niche for Prochlorococcus sp. in low-light suboxic environments in the Arabian Sea and the Eastern Tropical North Pacific. Deep Sea Research Part I: Oceanographic Research Papers 47: 1183–1205.

Goldstein, L., and D. Funkhouser. 1972. Biosynthesis of trimethylamine oxide in the nurse shark, *Ginglymostoma cirratum*. Comparative Biochemistry and Physiology Part A: Physiology 42: 51–57.

González-Rodríguez, E., A. Trasviña-Castro, G. Gaxiola-Castro, L. Zamudio, and R. Cervantes-Duarte. 2012. Net primary productivity, upwelling and coastal currents in the Gulf of Ulloa, Baja California, México. Ocean Science 8: 703–711.

Gordon, L., J. Jennings, A. Ross, and J. Krest. 1993. A Suggested Protocol for Continuous Flow Automated Analysis of Seawater Nutrients (Phosphate, Nitrate, Nitrite and Silicic Acid) in the WOCE Hydrographic Program and the Joint Global Ocean Fluxes Study. Methods Manual WHPO 91–1.

Hagemann, M. 2011. Molecular biology of cyanobacterial salt acclimation. FEMS Microbiology Reviews 35: 87–123.

Halsey, K. H., A. E. Carter, and S. J. Giovannoni. 2012. Synergistic metabolism of a broad range of C1 compounds in the marine methylotrophic bacterium HTCC2181. Environmental Microbiology 14: 630–640.

Haro-Moreno, J. M., F. Rodriguez-Valera, R. Rosselli, F. Martinez-Hernandez, J. J. Roda-Garcia, M. L. Gomez, O. Fornas, et al. 2019. Ecogenomics of the SAR11 clade. Environmental Microbiology 22: 1748.

Heal, K. R., B. P. Durham, A. K. Boysen, L. T. Carlson, W. Qin, F. Ribalet, A. E. White, et al. 2021. Marine Community Metabolomes Carry Fingerprints of Phytoplankton Community Composition. mSystems 6: e01334–20.

Heal, K. R., N. A. Kellogg, L. T. Carlson, R. M. Lionheart, and A. E. Ingalls. 2019. Metabolic Consequences of Cobalamin Scarcity in the Diatom *Thalassiosira pseudonana* as Revealed Through Metabolomics. Protist 170: 328–348.

Hyatt, D., G.-L. Chen, P. F. LoCascio, M. L. Land, F. W. Larimer, and L. J. Hauser. 2010. Prodigal: prokaryotic gene recognition and translation initiation site identification. BMC Bioinformatics 11: 119.

Jaffe, A. L., K. Harrison, R. Z. Wang, L. J. Taylor-Kearney, N. Jain, N. Prywes, P. M. Shih, et al. 2024. Cyanobacteria from marine oxygen-deficient zones encode both form I and form II Rubiscos. Proceedings of the National Academy of Sciences of the United States of America 121: e2418345121.

Johnson, W. M., M. C. Kido Soule, K. Longnecker, M. P. Bhatia, S. J. Hallam, M. W. Lomas, and E. B. Kujawinski. 2023. Particulate and dissolved metabolite distributions along a latitudinal transect of the western Atlantic Ocean. Limnology and Oceanography 68: 377–393.

Johnson, W. M., K. Longnecker, M. C. Kido Soule, W. A. Arnold, M. P. Bhatia, S. J. Hallam, B. A. S. Van Mooy, and E. B. Kujawinski. 2020. Metabolite composition of sinking particles differs from surface suspended particles across a latitudinal transect in the South Atlantic. Limnology and Oceanography 65: 111–127.

Johnson, Z., M. L. Landry, R. R. Bidigare, S. L. Brown, L. Campbell, J. Gunderson, J. Marra, and C. Trees. 1999. Energetics and growth kinetics of a deep *Prochlorococcus* spp. population in the Arabian Sea. Deep Sea Research Part II: Topical Studies in Oceanography 46: 1719–1743.

Karl, D. M., K. M. Björkman, M. J. Church, L. A. Fujieki, E. M. Grabowski, and R. M. Letelier. 2022. Temporal dynamics of total microbial biomass and particulate detritus at Station ALOHA. Progress in Oceanography 205: 102803.

Keeling, P. J., F. Burki, H. M. Wilcox, B. Allam, E. E. Allen, L. A. Amaral-Zettler, E. V. Armbrust, et al. 2014. The Marine Microbial Eukaryote Transcriptome Sequencing Project (MMETSP): Illuminating the Functional Diversity of Eukaryotic Life in the Oceans through Transcriptome Sequencing. PLoS Biology 12: e1001889.

Keller, M., W. Bellows, and R. Guillard. 1989. Dimethyl Sulfide Production in Marine Phytoplankton. Biogenic Sulfur in the Environment 393.

Kessner, D., M. Chambers, R. Burke, D. Agus, and P. Mallick. 2008. ProteoWizard: open source software for rapid proteomics tools development. Bioinformatics 24: 2534–2536.

Kettler, G. C., A. C. Martiny, K. Huang, J. Zucker, M. L. Coleman, S. Rodrigue, F. Chen, et al. 2007. Patterns and Implications of Gene Gain and Loss in the Evolution of Prochlorococcus. PLOS Genetics 3: e231.

Kettles, N. L., S. Kopriva, and G. Malin. 2014. Insights into the Regulation of DMSP Synthesis in the Diatom Thalassiosira pseudonana through APR Activity, Proteomics and Gene Expression Analyses on Cells Acclimating to Changes in Salinity, Light and Nitrogen. PLOS ONE 9: e94795.

Kiko, R., H. Hauss, F. Buchholz, and F. Melzner. 2016. Ammonium excretion and oxygen respiration of tropical copepods and euphausiids exposed to oxygen minimum zone conditions. Biogeosciences 13: 2241–2255.

Klähn, S., and M. Hagemann. 2011. Compatible solute biosynthesis in cyanobacteria. Environmental Microbiology 13: 551–562.

Klähn, S., C. Steglich, W. R. Hess, and M. Hagemann. 2010. Glucosylglycerate: a secondary compatible solute common to marine cyanobacteria from nitrogen-poor environments. Environmental Microbiology 12: 83–94.

Klemetsen, T., I. A. Raknes, J. Fu, A. Agafonov, S. V. Balasundaram, G. Tartari, E. Robertsen, and N. P. Willassen. 2018. The MAR databases: development and implementation of databases specific for marine metagenomics. Nucleic Acids Research 46: D692–D699.

Kopylova, E., L. Noé, and H. Touzet. 2012. SortMeRNA: fast and accurate filtering of ribosomal RNAs in metatranscriptomic data. Bioinformatics 28: 3211–3217.

Kujawinski, E. B., R. Braakman, K. Longnecker, J. W. Becker, S. W. Chisholm, K. Dooley, M. C. Kido Soule, et al. 2023. Metabolite diversity among representatives of divergent Prochlorococcus ecotypes. mSystems 8: e01261–22.

Lai, M.-C., D.-R. Yang, and M.-J. Chuang. 1999. Regulatory Factors Associated with Synthesis of the Osmolyte Glycine Betaine in the Halophilic Methanoarchaeon Methanohalophilus portucalensis. Applied and Environmental Microbiology 65: 828.

Langmead, B., and S. L. Salzberg. 2012. Fast gapped-read alignment with Bowtie 2. Nature Methods 9: 357–359.

Lavin, P., B. González, J. F. Santibáñez, D. J. Scanlan, and O. Ulloa. 2010. Novel lineages of Prochlorococcus thrive within the oxygen minimum zone of the eastern tropical South Pacific. Environmental Microbiology Reports 2: 728–738.

Li, C.-Y., X.-L. Chen, D. Zhang, P. Wang, Q. Sheng, M. Peng, B.-B. Xie, et al. 2017. Structural mechanism for bacterial oxidation of oceanic trimethylamine into trimethylamine N-oxide. Molecular Microbiology 103: 992–1003.

Li, D., C.-M. Liu, R. Luo, K. Sadakane, and T.-W. Lam. 2015. MEGAHIT: an ultra-fast single-node solution for large and complex metagenomics assembly via succinct de Bruijn graph. Bioinformatics 31: 1674–1676.

Lidbury, I. D. E. A., J. C. Murrell, and Y. Chen. 2015. Trimethylamine and trimethylamine N-oxide are supplementary energy sources for a marine heterotrophic bacterium: implications for marine carbon and nitrogen cycling. The ISME Journal 9: 760–769.

Lidbury, I., J. C. Murrell, and Y. Chen. 2014. Trimethylamine N-oxide metabolism by abundant marine heterotrophic bacteria. Proceedings of the National Academy of Sciences 111: 2710–2715.

Longnecker, K., M. C. Kido Soule, G. J. Swarr, R. J. Parsons, S. Liu, W. M. Johnson, B. Widner, et al. 2024. Seasonal and daily patterns in known dissolved metabolites in the northwestern Sargasso Sea. Limnology and Oceanography 69: 449–466.

Lu, W.-D., Z.-M. Chi, and C.-D. Su. 2006. Identification of glycine betaine as compatible solute in Synechococcus sp. WH8102 and characterization of its N-methyltransferase genes involved in betaine synthesis. Archives of Microbiology 186: 495–506.

Maas, A. E., S. L. Frazar, D. M. Outram, B. A. Seibel, and K. F. Wishner. 2014. Fine-scale vertical distribution of macroplankton and micronekton in the Eastern Tropical North Pacific in association with an oxygen minimum zone. Journal of Plankton Research 36: 1557–1575.

Maas, A. E., K. F. Wishner, and B. A. Seibel. 2012. Metabolic suppression in thecosomatous pteropods as an effect of low temperature and hypoxia in the eastern tropical North Pacific. Marine Biology 159: 1955–1967.

Maker, J. R., A. Struempler, and S. Chaykin. 1963. A comparative study of trimethylamine-*N*-oxide biosynthesis. Biochimica et Biophysica Acta 71: 58–64.

Mattes, T. E., S. Burke, G. Rocap, and R. M. Morris. 2022. Two Metatranscriptomic Profiles through Low-Dissolved-Oxygen Waters (DO, 0 to 33 µM) in the Eastern Tropical North Pacific Ocean. Microbiology Resource Announcements 11: e01201–21.

Mausz, M. A., R. L. Airs, J. L. Dixon, C. E. Widdicombe, G. A. Tarran, L. Polimene, S. Dashfield, et al. 2022. Microbial uptake dynamics of choline and glycine betaine in coastal seawater. Limnology and Oceanography 67: 1052–1064.

Measures, J. C. 1975. Role of amino acids in osmoregulation of non-halophilic bacteria. Nature 257: 398–400.

Menzel, P., K. L. Ng, and A. Krogh. 2016. Fast and sensitive taxonomic classification for metagenomics with Kaiju. Nature Communications 7: 11257.

Moran, M. A., E. B. Kujawinski, W. F. Schroer, S. A. Amin, N. R. Bates, E. M. Bertrand, R. Braakman, et al. 2022. Microbial metabolites in the marine carbon cycle. Nature Microbiology 7: 508–523.

Morris, R. M., M. S. Rappé, S. A. Connon, K. L. Vergin, W. A. Siebold, C. A. Carlson, and S. J. Giovannoni. 2002. SAR11 clade dominates ocean surface bacterioplankton communities. Nature 420: 806–810.

Nakamura, H., K. Fujimaki, O. Sampei, and A. Murai. 1993. Gonyol: Methionine-induced sulfonium accumulation in a dinoflagellate Gonyaulax polyedra. Tetrahedron Letters 34: 8481–8484.

Narváez-Barragán, D. A., M. Sperfeld, and E. Segev. 2024. The bacterial Bmt methionine synthase is involved in lag phase shortening. 2024.06.19.599700.

Nguyen, L.-T., H. A. Schmidt, A. von Haeseler, and B. Q. Minh. 2015. IQ-TREE: A Fast and Effective Stochastic Algorithm for Estimating Maximum-Likelihood Phylogenies. Molecular Biology and Evolution 32: 268–274.

Noell, S. E., and S. J. Giovannoni. 2019. SAR11 bacteria have a high affinity and multifunctional glycine betaine transporter. Environmental Microbiology 21: 2559–2575.

Nyyssölä, A., J. Kerovuo, P. Kaukinen, N. Von Weymarn, and T. Reinikainen. 2000. Extreme Halophiles Synthesize Betaine from Glycine by Methylation. Journal of Biological Chemistry 275: 22196–22201.

O’Leary, N. A., M. W. Wright, J. R. Brister, S. Ciufo, D. Haddad, R. McVeigh, B. Rajput, et al. 2016. Reference sequence (RefSeq) database at NCBI: current status, taxonomic expansion, and functional annotation. Nucleic Acids Research 44: D733–745.

Parris, D. J., S. Ganesh, V. P. Edgcomb, E. F. DeLong, and F. J. Stewart. 2014. Microbial eukaryote diversity in the marine oxygen minimum zone off northern Chile. Frontiers in Microbiology 5.

Poretsky, R. S., S. Sun, X. Mou, and M. A. Moran. 2010. Transporter genes expressed by coastal bacterioplankton in response to dissolved organic carbon. Environmental Microbiology 12: 616–627.

Rocap, G., F. W. Larimer, J. Lamerdin, S. Malfatti, P. Chain, N. A. Ahlgren, A. Arellano, et al. 2003. Genome divergence in two Prochlorococcus ecotypes reflects oceanic niche differentiation. Nature 424: 1042–1047.

Ruiz-Perez, C. A., A. D. Bertagnolli, D. Tsementzi, T. Woyke, F. J. Stewart, and K. T. Konstantinidis. 2021. Description of *Candidatus* Mesopelagibacter carboxydoxydans and *Candidatus* Anoxipelagibacter denitrificans: Nitrate-reducing SAR11 genera that dominate mesopelagic and anoxic marine zones. Systematic and Applied Microbiology 44: 126185.

Satinsky, B. M., S.M. Gifford, B. C. Crump and M. A. Moran. 2013. Use of internal standards for quantitative metatranscriptome and metagenome analysis. Meth Enzymol, ed DeLong EF (Academic, Burlington, MA), Vol 531, pp 237–250.

Scanlan, D. J., M. Ostrowski, S. Mazard, A. Dufresne, L. Garczarek, W. R. Hess, A. F. Post, et al. 2009. Ecological Genomics of Marine Picocyanobacteria. Microbiology and Molecular Biology Reviews 73: 249–299.

Seibel, B. A., and P. J. Walsh. 2002. Trimethylamine oxide accumulation in marine animals: relationship to acylglycerol storagej. Journal of Experimental Biology 205: 297–306.

Strom, A. R. 1979. Biosynthesis of trimethylamine oxide in calanoid copepods. seasonal changes in trimethylamine monooxygenase activity. Marine Biology 51: 33–40.

Sud, M., E. Fahy, D. Cotter, K. Azam, I. Vadivelu, C. Burant, A. Edison, et al. 2016. Metabolomics Workbench: An international repository for metabolomics data and metadata, metabolite standards, protocols, tutorials and training, and analysis tools. Nucleic Acids Research 44: D463–D470.

Sun, J., M. A. Mausz, Y. Chen, and S. J. Giovannoni. 2019. Microbial trimethylamine metabolism in marine environments. Environmental Microbiology 21: 513–520.

Sun, J., L. Steindler, J. C. Thrash, K. H. Halsey, D. P. Smith, A. E. Carter, Z. C. Landry, and S. J. Giovannoni. 2011. One Carbon Metabolism in SAR11 Pelagic Marine Bacteria. PLOS ONE 6: e23973.

Tiano, L., E. Garcia-Robledo, T. Dalsgaard, A. H. Devol, B. B. Ward, O. Ulloa, D. E. Canfield, and N. Peter Revsbech. 2014. Oxygen distribution and aerobic respiration in the north and south eastern tropical Pacific oxygen minimum zones. Deep Sea Research Part I: Oceanographic Research Papers 94: 173–183.

Treberg, J. R., and W. R. Driedzic. 2002. Elevated levels of trimethylamine oxide in deep-sea fish: evidence for synthesis and intertissue physiological importance. Journal of Experimental Zoology 293: 39–45.

Tripp, H. J. 2013. The unique metabolism of SAR11 aquatic bacteria. Journal of Microbiology 51: 147–153.

Tsementzi, D., J. Wu, S. Deutsch, S. Nath, L. M. Rodriguez-R, A. S. Burns, P. Ranjan, et al. 2016. SAR11 bacteria linked to ocean anoxia and nitrogen loss. Nature 536: 179–183.

Ulloa, O., C. Henríquez-Castillo, S. Ramírez-Flandes, A. M. Plominsky, A. A. Murillo, C. Morgan-Lang, S. J. Hallam, and R. Stepanauskas. 2021. The cyanobacterium Prochlorococcus has divergent light-harvesting antennae and may have evolved in a low-oxygen ocean. Proceedings of the National Academy of Sciences of the United States of America 118: e2025638118.

Vorobev, A., S. Sharma, M. Yu, J. Lee, B. J. Washington, W. B. Whitman, F. Ballantyne IV, et al. 2018. Identifying labile DOM components in a coastal ocean through depleted bacterial transcripts and chemical signals. Environmental Microbiology 20: 3012–3030.

Waditee, R., Y. Tanaka, K. Aoki, T. Hibino, H. Jikuya, J. Takano, T. Takabe, and T. Takabe. 2003. Isolation and functional characterization of N-methyltransferases that catalyze betaine synthesis from glycine in a halotolerant photosynthetic organism Aphanothece halophytica. The Journal of Biological Chemistry 278: 4932–4942.

Wakeham, S. G. 2020. Organic biogeochemistry in the oxygen-deficient ocean: A review. Organic Geochemistry 149: 104096.

Welsh, D. T. 2000. Ecological significance of compatible solute accumulation by micro-organisms: from single cells to global climate. FEMS Microbiology Reviews 24: 263–290.

Widner, B., C. A. Fuchsman, B. X. Chang, G. Rocap, and M. R. Mulholland. 2018. Utilization of urea and cyanate in waters overlying and within the eastern tropical north Pacific oxygen deficient zone. FEMS Microbiology Ecology 94: fiy138.

Wishner, K. F., B. A. Seibel, C. Roman, C. Deutsch, D. Outram, C. T. Shaw, M. A. Birk, et al. 2018. Ocean deoxygenation and zooplankton: Very small oxygen differences matter. Science Advances 4: eaau5180.

Wong, J. C. Y., J. A. Raven, M. Aldunate, S. Silva, J. D. Gaitán-Espitia, C. A. Vargas, O. Ulloa, and P. von Dassow. 2023. Do phytoplankton require oxygen to survive? A hypothesis and model synthesis from oxygen minimum zones. Limnology and Oceanography 68: 1417–1437.

Xu, Z., S. Cheung, H. Endo, X. Xia, W. Wu, B. Chen, N. H. E. Ho, et al. 2022. Disentangling the Ecological Processes Shaping the Latitudinal Pattern of Phytoplankton Communities in the Pacific Ocean. mSystems 7: e01203–21.

Yang, N., R. Ding, and J. Liu. 2022. Synthesizing glycine betaine via choline oxidation pathway as an osmoprotectant strategy in Haloferacales. Gene 847: 146886.

Zheng, Y., J. Wang, S. Zhou, Y. Zhang, J. Liu, C.-X. Xue, B. T. Williams, et al. 2020. Bacteria are important dimethylsulfoniopropionate producers in marine aphotic and high-pressure environments. Nature Communications 11: 4658.

