## Supplemental Text for "Marine community metabolomes in the eastern tropical North Pacific Oxygen Deficient Zone reveal glycine betaine as a metabolic link between *Prochlorococcus* and SAR11"

### *Metabolite extraction*

Metabolites were extracted following the modified Bligh-Dyer protocol described in Boysen et al., 2018. Briefly, frozen filters were cut into small pieces and transferred to bead-beating tubes containing silica beads, cold aqueous solvent (50:50 methanol:water), and dichloromethane (DCM). Samples were homogenized using a FastPrep-24 for 30 seconds, incubated at –20 °C, and this cycle was repeated three times. After centrifugation (5,000 rpm, 90 seconds, 4 °C), the aqueous phase was collected and the remaining pellet was rinsed three times with 50:50 methanol:water. All aqueous layers were pooled and dried under nitrogen gas. The organic layer was similarly transferred, rinsed twice with cold DCM, centrifuged, and dried under nitrogen. Aqueous and organic fractions were reconstituted in 400 µL of water and 400 µL of 1:1 water:acetonitrile, respectively. Isotope-labeled internal standards were then added at the same concentrations as in Boysen et al., 2018. Only the aqueous fraction was analyzed in this study.

### *Liquid chromatography-mass spectrometry analysis*

### Reconstituted extracts were analyzed using Liquid chromatography-mass spectrometry (LC-MS) using both reversed-phase (RP) and hydrophilic interaction chromatography (HILIC), employing the same solvent systems, gradients, and columns as Boysen et al. (2018), a Waters Acquity UPLC HSS Cyano column (RP) and a SeQuant ZIC-pHILIC column (HILIC). Injection volumes were 5 µL for RP and 2 µL for HILIC analyses. Both methods were run on a Thermo Q-Exactive (QE) mass spectrometer with electrospray ionization (ESI). RP was operated in positive mode with 120,000 resolution, while HILIC used polarity switching with 60,000 resolution. Raw data files were converted to .mzXML using ProteoWizard, and peaks were manually integrated using Skyline. In-house quality control filters excluded peaks with a signal-to-noise ratio <4, peaks not three times above the seawater blank, and compounds not detected in at least three samples

*Data normalization*

Normalization was performed using the Best-Matched Internal Standard (B-MIS) method (Boysen et al., 2018). This method accounts for non-biological variability introduced by ion suppression, matrix effects, and analytical drift by selecting an internal standard for each compound that minimizes technical variability across pooled quality control (QC) samples. Full and half-strength pooled samples (1:1 dilution with water) were included throughout each batch to train the normalization algorithm.

Normalization was performed on a compound-specific basis. B-MIS normalization was used only for metabolites without a corresponding isotope-labeled internal standard. For metabolites with labeled analogs, quantification was based solely on the peak area ratio of analyte to internal standard, and B-MIS was not applied. For B-MIS compounds, each metabolite was evaluated against all candidate internal standards, and peak areas were divided by the signal of the internal standard normalized to its mean across pooled QC injections. If this correction reduced the relative standard deviation (RSD) by > 10%, the internal standard was assigned as the best match. Metabolites with low technical variability (RSD < 1%) were not normalized, as correction could introduce unnecessary noise. This approach enabled targeted correction of obscuring variation while preserving high-confidence raw measurements where appropriate.

*Metabolite Quantification*

Quantification followed previously established approaches (Boysen et al., 2021; Heal et al., 2021). For analytes with isotope-labeled standards, concentrations were calculated using:

$${Concentration}_{M}= \frac{{Area}_{M}}{{Area}_{IS}}*\left[ IS \right]_{vial}* \frac{V_{reconst}}{V_{filtered}}$$

Where M = metabolite, IS = internal standard, Area = peak area, and V = volume.

For other compounds, absolute concentrations were estimated using:

$${Concentration}_{M}= \frac{{Area}_{M}}{RF}* \frac{V_{reconst}}{V_{filtered}} * \frac{1}{{RF}_{ratio}}$$

where RF is the response factor (area/concentration) in water and RF_ratio_ is the ratio of RF in matrix to RF in water, as described in the work of Boysen et al., 2018. RF values were determined per compound within each LC-MS batch for compounds lacking paired isotope-labeled standards. This approach provides a reproducible basis for relative quantification, as demonstrated in previous work using complex environmental matrices. Volume filtered were ~10 L and volume reconstituted were 400 µl for each sample

For the compound homarine, which was later found to be contaminated in the commercial standard (Santa Cruz Biotechnology, lot B1916), concentrations were corrected using newly acquired isotope-labeled standards, reducing the originally reported values by a factor of 18.

All concentration data are reported in units of nanomoles per liter (nM) and are corrected for extraction volume and matrix effects. B-MIS normalization and quantification were implemented in R using custom scripts modified from those provided in Boysen et al., 2018 available at <https://github.com/IngallsLabUW/B-MIS-normalization>.

**Citations**

Boysen, A. K., L. T. Carlson, B. P. Durham, R. D. Groussman, F. O. Aylward, F. Ribalet, K. R. Heal, et al. 2021. Particulate Metabolites and Transcripts Reflect Diel Oscillations of Microbial Activity in the Surface Ocean. *mSystems* 6: 10.1128/msystems.00896-20.

Boysen, A. K., K. R. Heal, L. T. Carlson, and A. E. Ingalls. 2018. Best-Matched Internal Standard Normalization in Liquid Chromatography–Mass Spectrometry Metabolomics Applied to Environmental Samples. *Analytical Chemistry* 90: 1363–1369.

Heal, K. R., B. P. Durham, A. K. Boysen, L. T. Carlson, W. Qin, F. Ribalet, A. E. White, et al. 2021. Marine Community Metabolomes Carry Fingerprints of Phytoplankton Community Composition. *mSystems* 6: e01334-20.
