## Supplemental Figures for "Marine community metabolomes in the eastern tropical North Pacific Oxygen Deficient Zone reveal glycine betaine as a metabolic link between *Prochlorococcus* and SAR11"

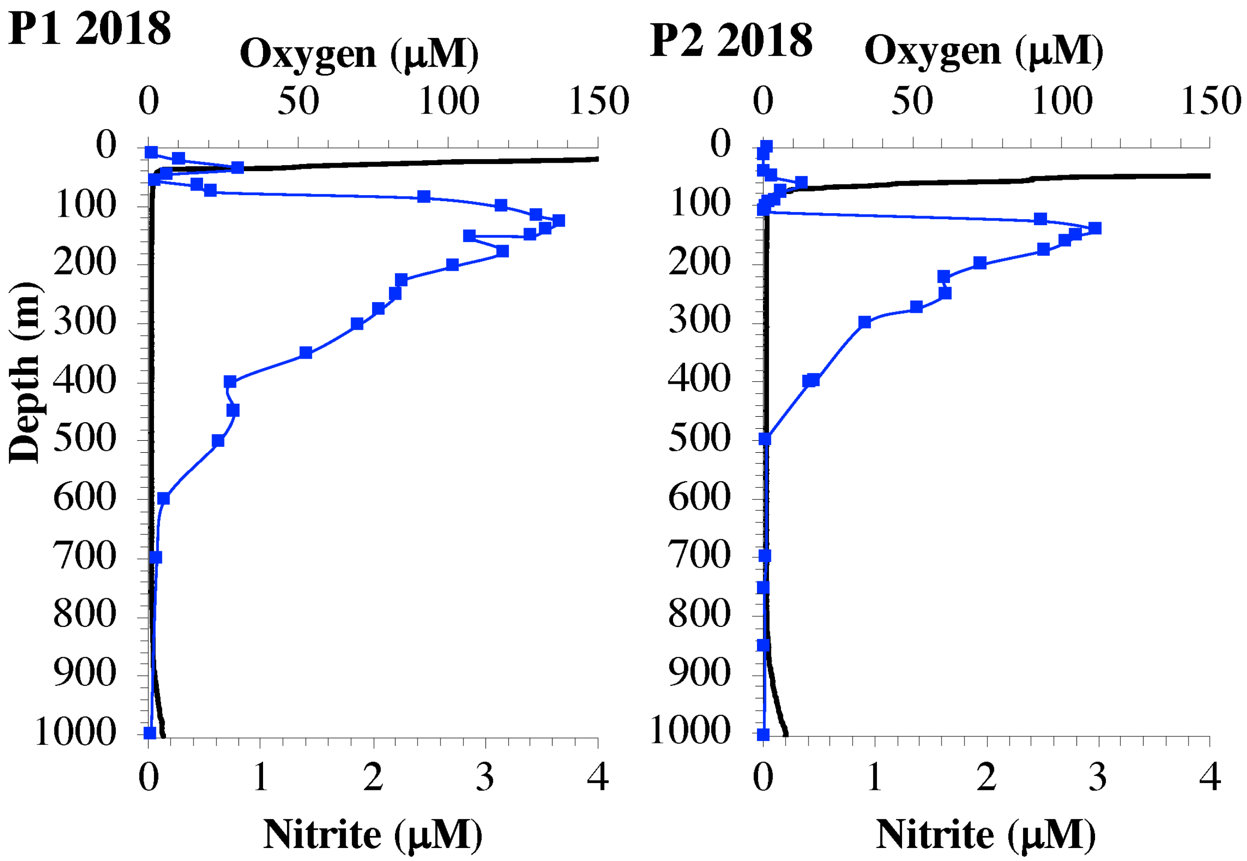


**Figure S1:** Depth profiles of nitrite (blue) and CTD measurements of oxygen (black) at the coastal ODZ (St. P1) offshore ODZ (St. P2).

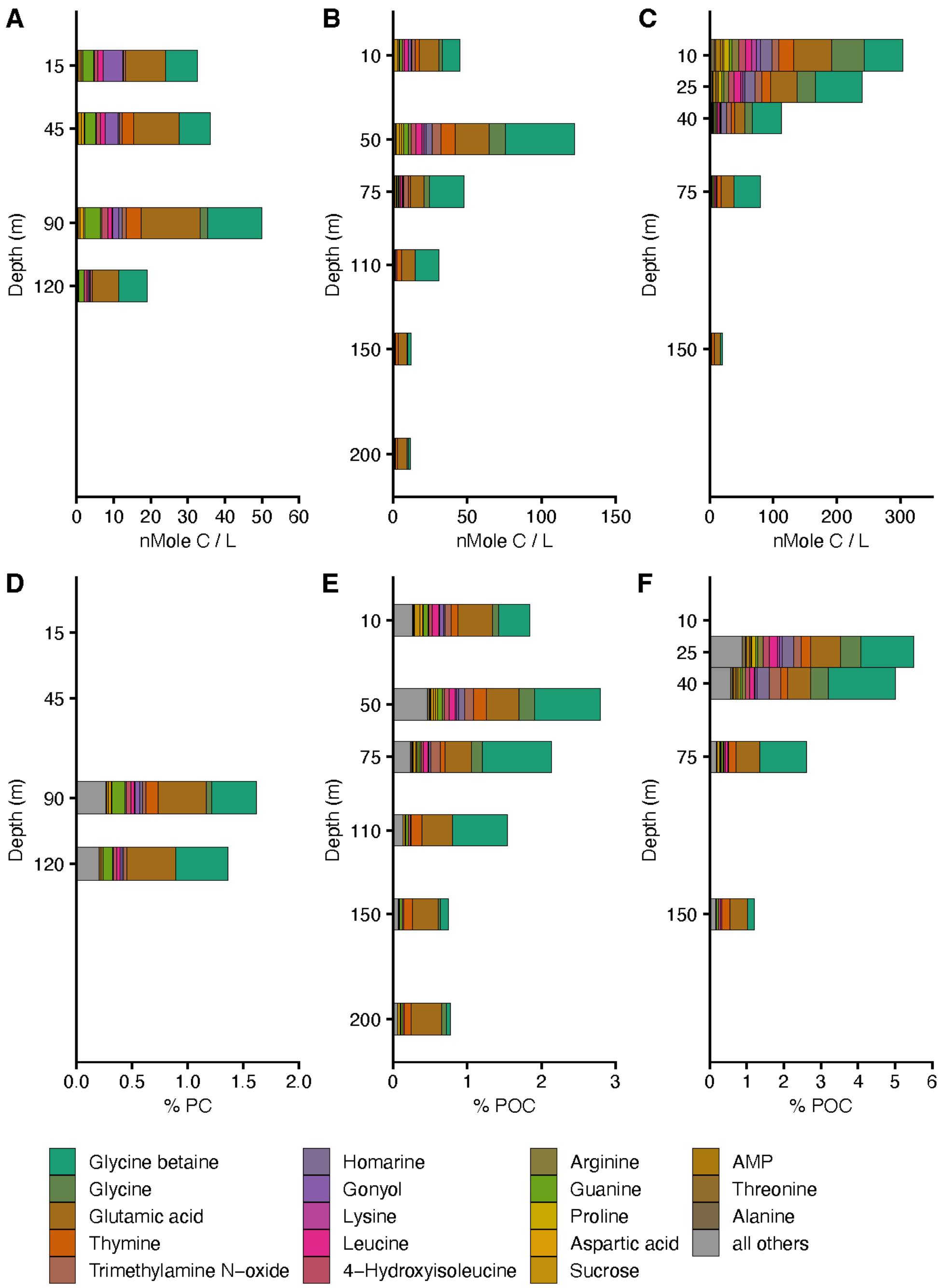


**Figure S2:** Most abundant 18 metabolites in environmental samples, presented as nmol carbon per L from the oxic NPG (**A**), offshore ODZ (St. P2) (**B**), and coastal ODZ (St. P1) (**C**) and percent particulate carbon from the oxic NPG (**D**), offshore ODZ (St. P2) (**E**), and coastal ODZ (St. P1) (**F**).

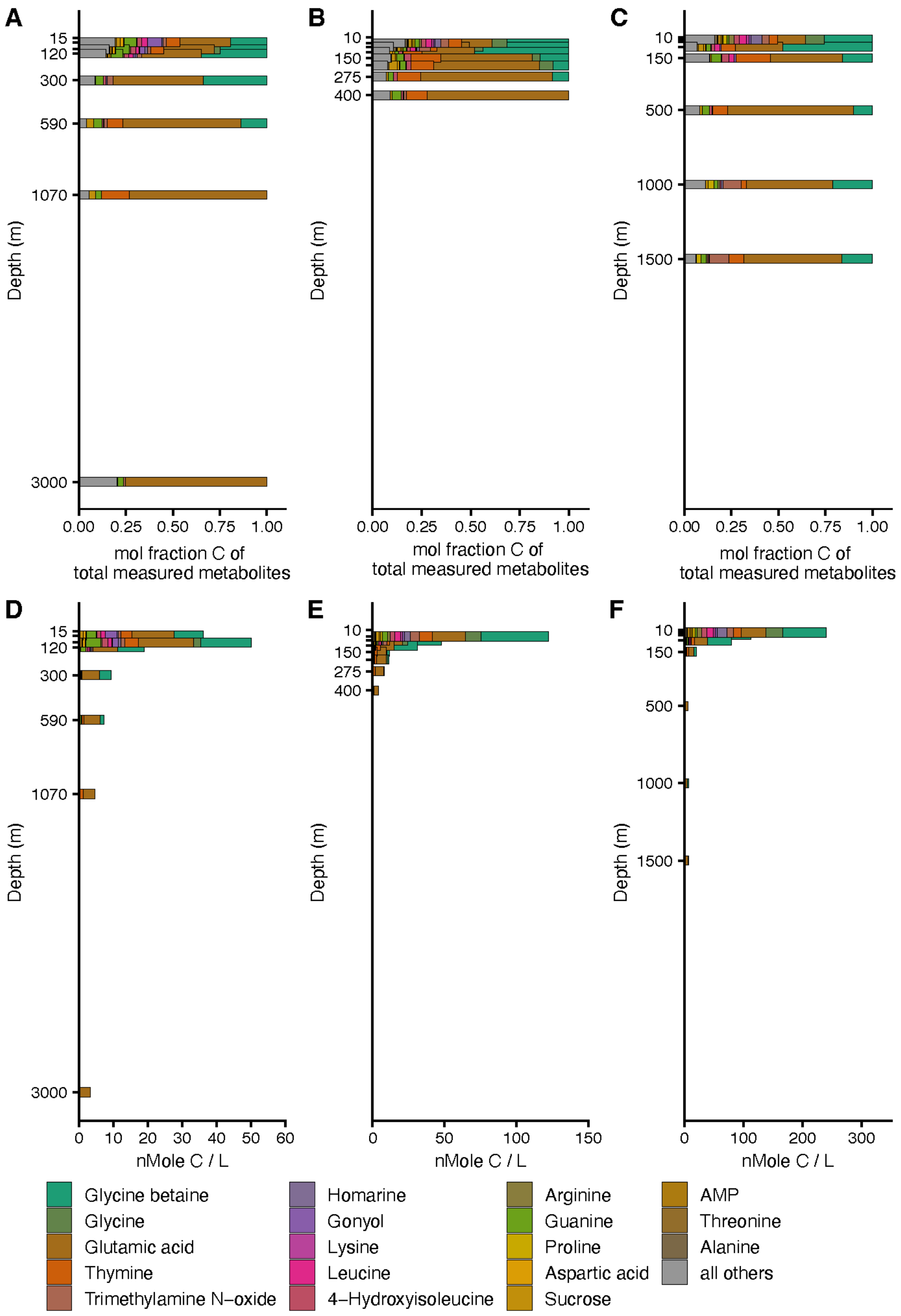


**Figure S3:** Depth profiles of the 18 most abundant metabolites across all environmental samples, including depths below 300 meters. Data are presented as either (i) mole fraction carbon, consistent with Figure 2, or (ii) total measured metabolites as nmol carbon per liter, consistent with Figure S2. Profiles as mole fraction carbon are shown for the oxic NPG **(A)**, the offshore ODZ (St. P2) (**B**), and the coastal ODZ (St. P1) (**C**). Additional metabolite profiles in nmol carbon per liter are shown for the oxic NPG (**D**), the offshore ODZ (St. P2) (**E**), and the coastal ODZ (St. P1) (**F**).

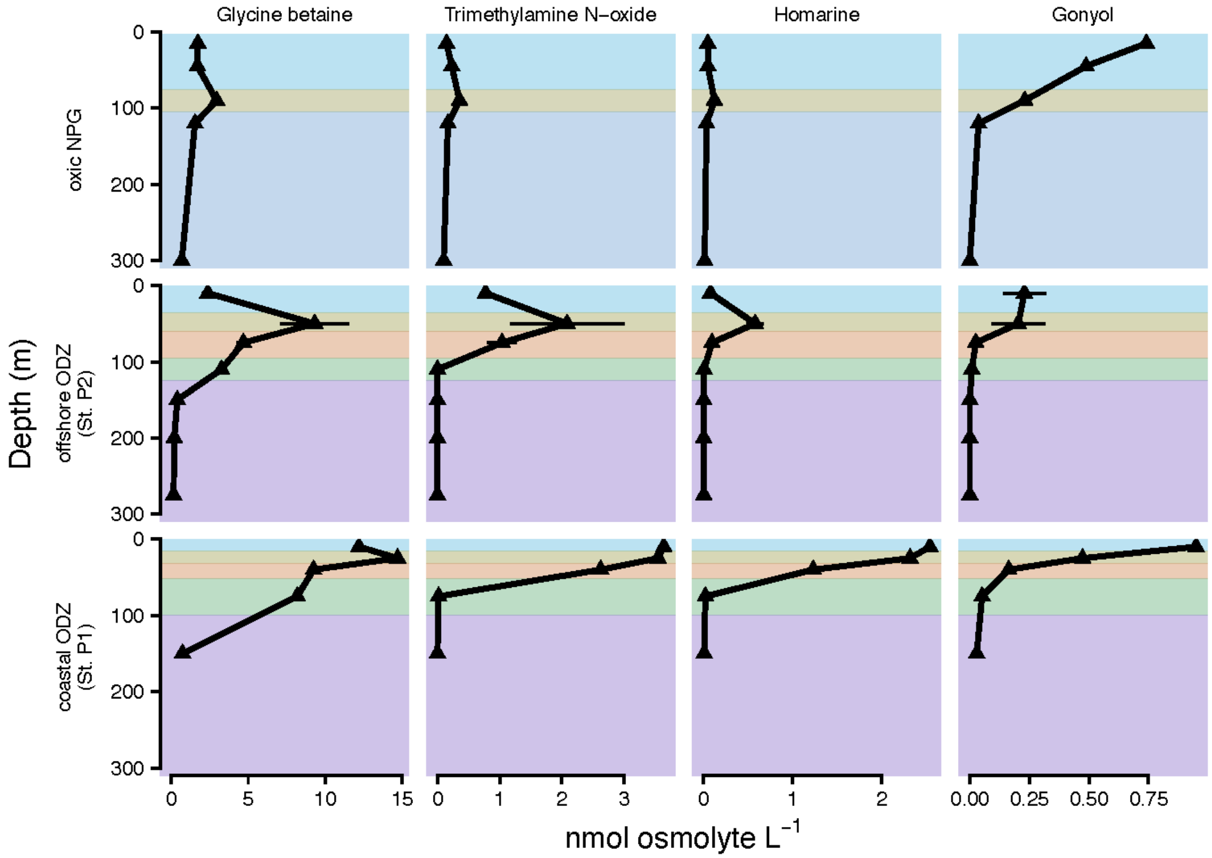


**Figure S4:** Depth profiles of the most abundant osmolytes, measured in nmol per L, within the upper 300 meters at each station. The metabolites shown, from left to right, are glycine betaine, trimethylamine N-oxide (TMAO), homarine, and gonyol.

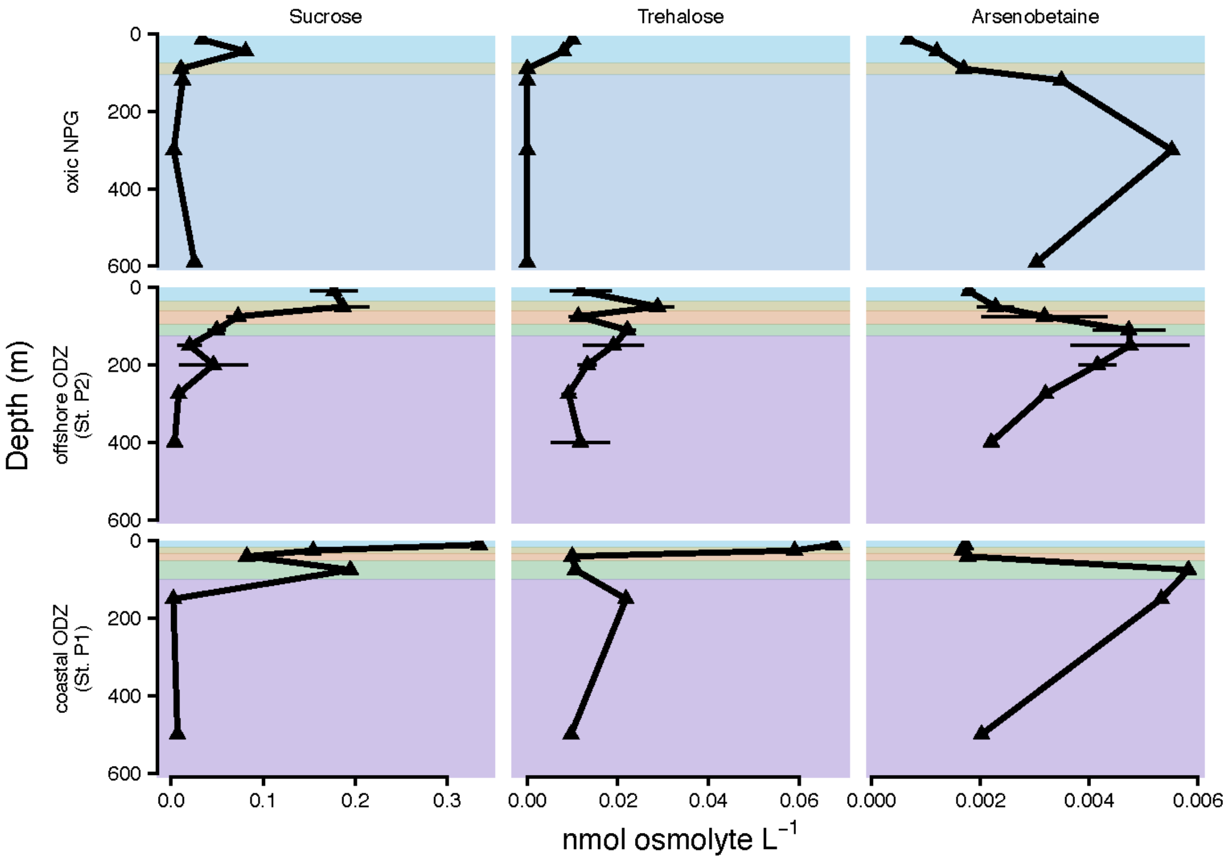


**Figure S5:** Depth profiles of some of the most abundant osmolytes, measured in nmol per liter, within the upper 600 meters at each station. The metabolites shown, from left to right, are sucrose, trehalose, and arsenobetaine.

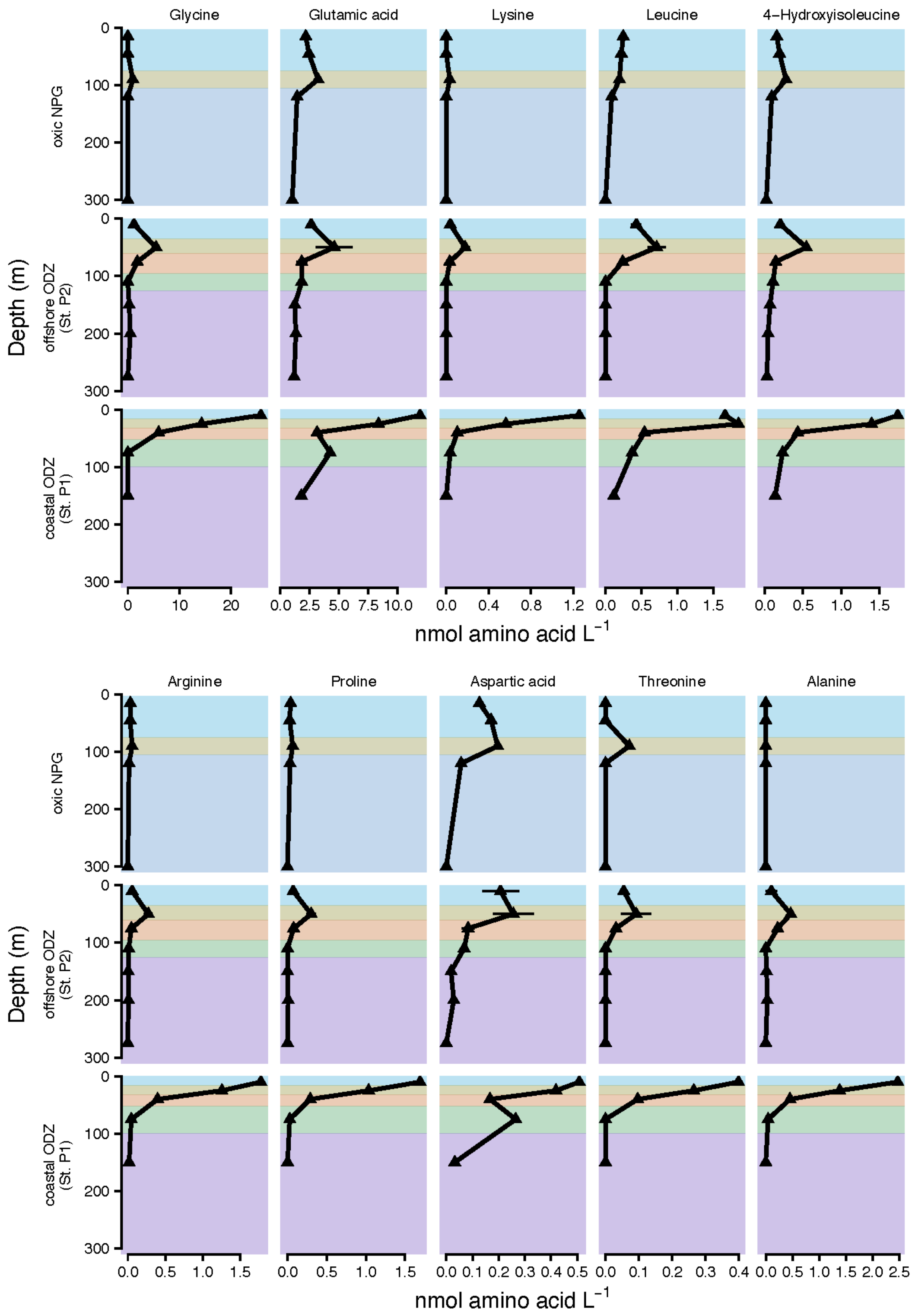


**Figure S6:** Depth profiles of the most abundant amino acids, measured in nmol per L, within the upper 300 meters at each station. The metabolites shown in the top row, from left to right, are glycine, glutamic acid, lysine, leucine, and 4-hydroxyisoleucine. The metabolites shown in the bottom row, from left to right, are arginine, proline, aspartic acid, threonine, and alanine

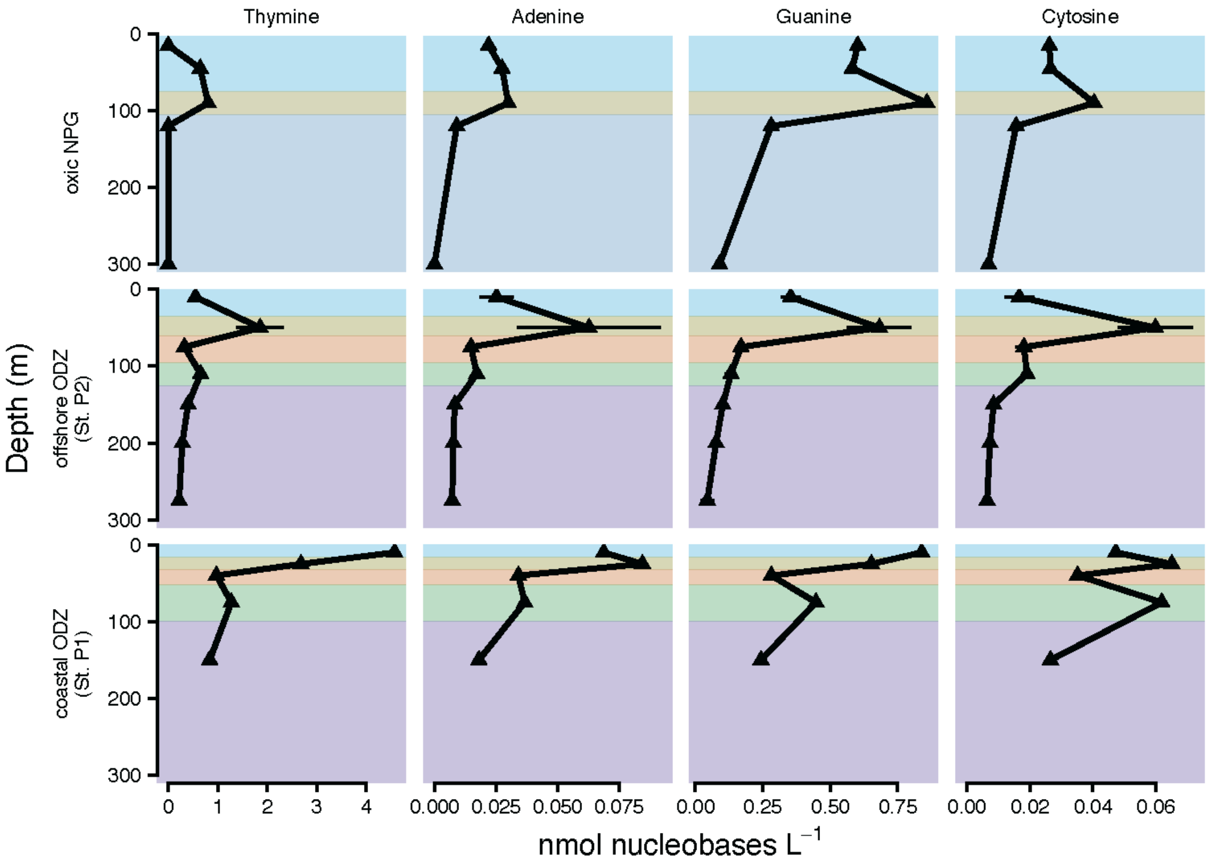


**Figure S7:** Depth profiles of nucleobases, measured in nmol per liter, within the upper 300 meters at each station. The metabolites shown, from left to right, are thymine, adenine, guanine, and cytosine.

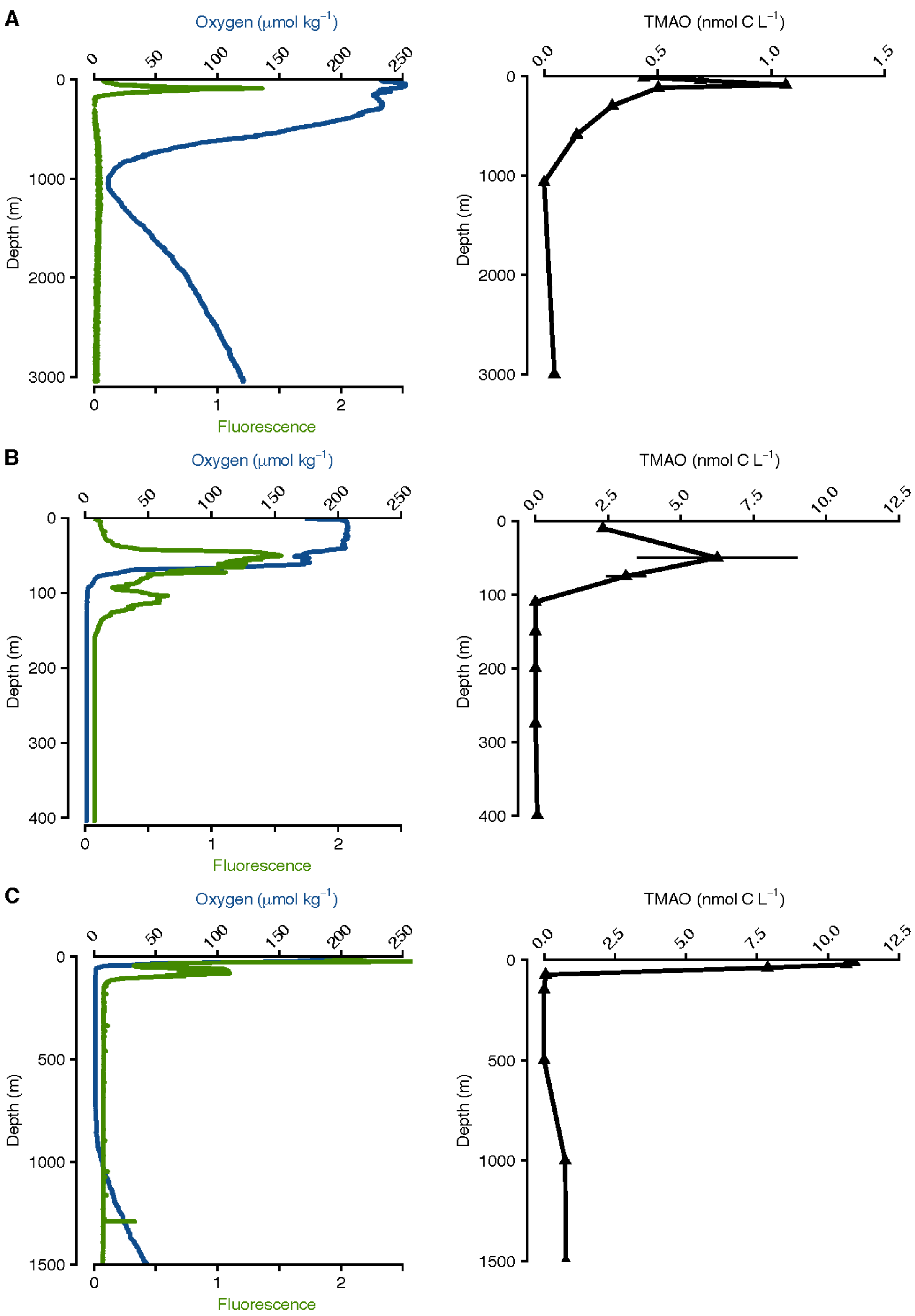


**Figure S8:** Trimethylamine N-oxide (TMAO) depth profiles as concentration of carbon and CTD measurements of oxygen and fluorescence. Depth profiles from oxic NPG **(A)**, offshore ODZ (St. P2) **(B)**, and coastal ODZ (St. P1) **(C)**. Locations of samples are shown in Figure 1.

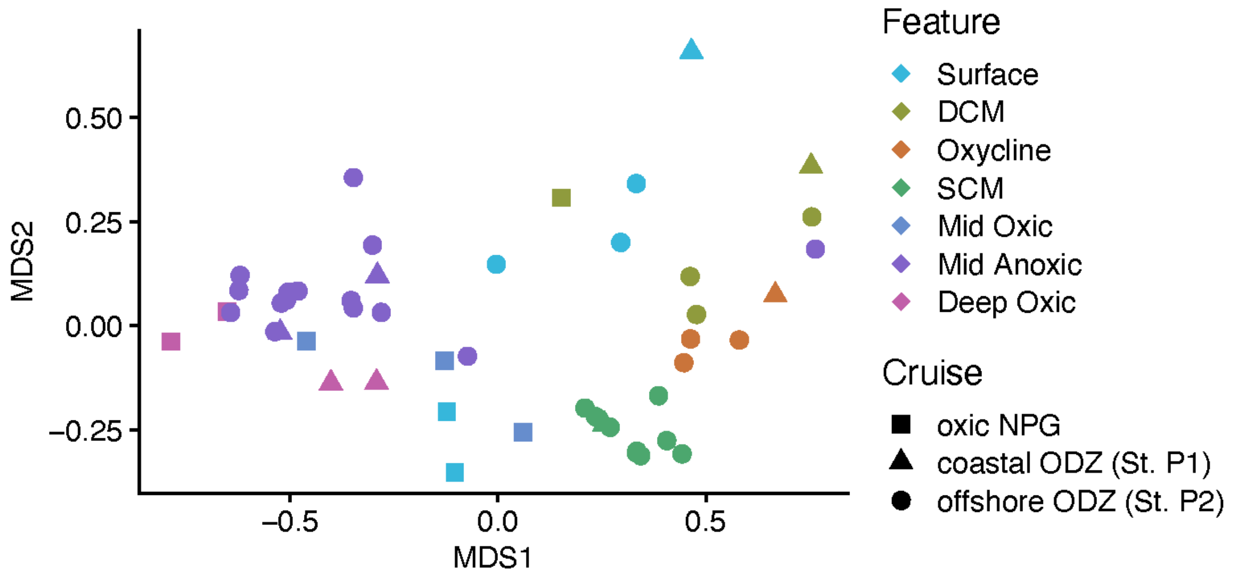


**Figure S9:** Nonmetric multidimensional scaling (NMDS) comparison of metabolite composition of three depth profiles (stress = 0.0887, $p < 0.01$ by Monte Carlo permutation). Colors are of zones based on environmental parameters in this study. This is based on Euclidean distance of quantified metabolite concentrations, normalized to the maximum value for each metabolite.

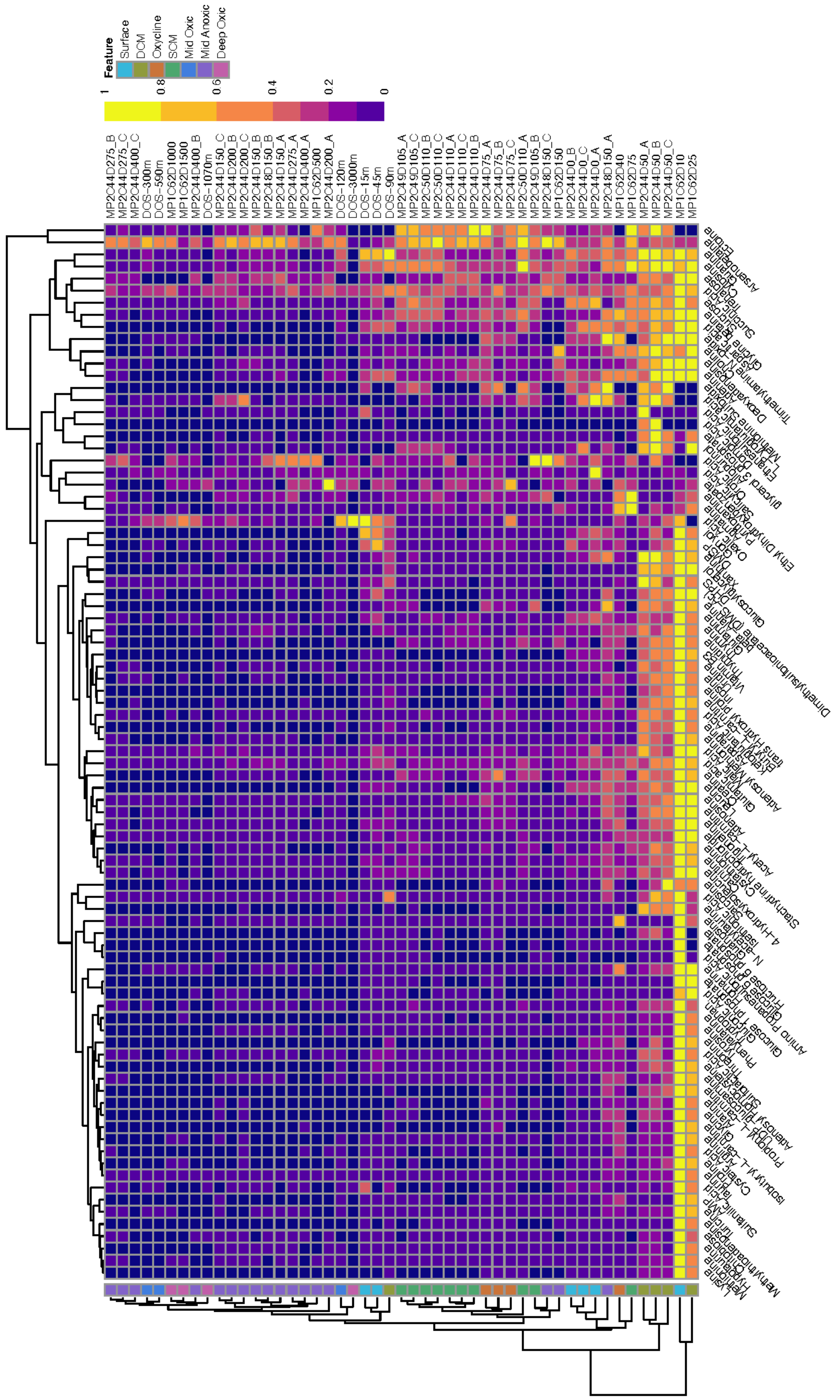


**Figure S10:** Hierarchical clustering dendrogram based on Euclidean distance of quantified metabolite concentrations, normalized to the maximum value for each metabolite. The accompanying heatmap displays the maximum normalized abundance for 87 total metabolites. See Table S1 for environmental context.

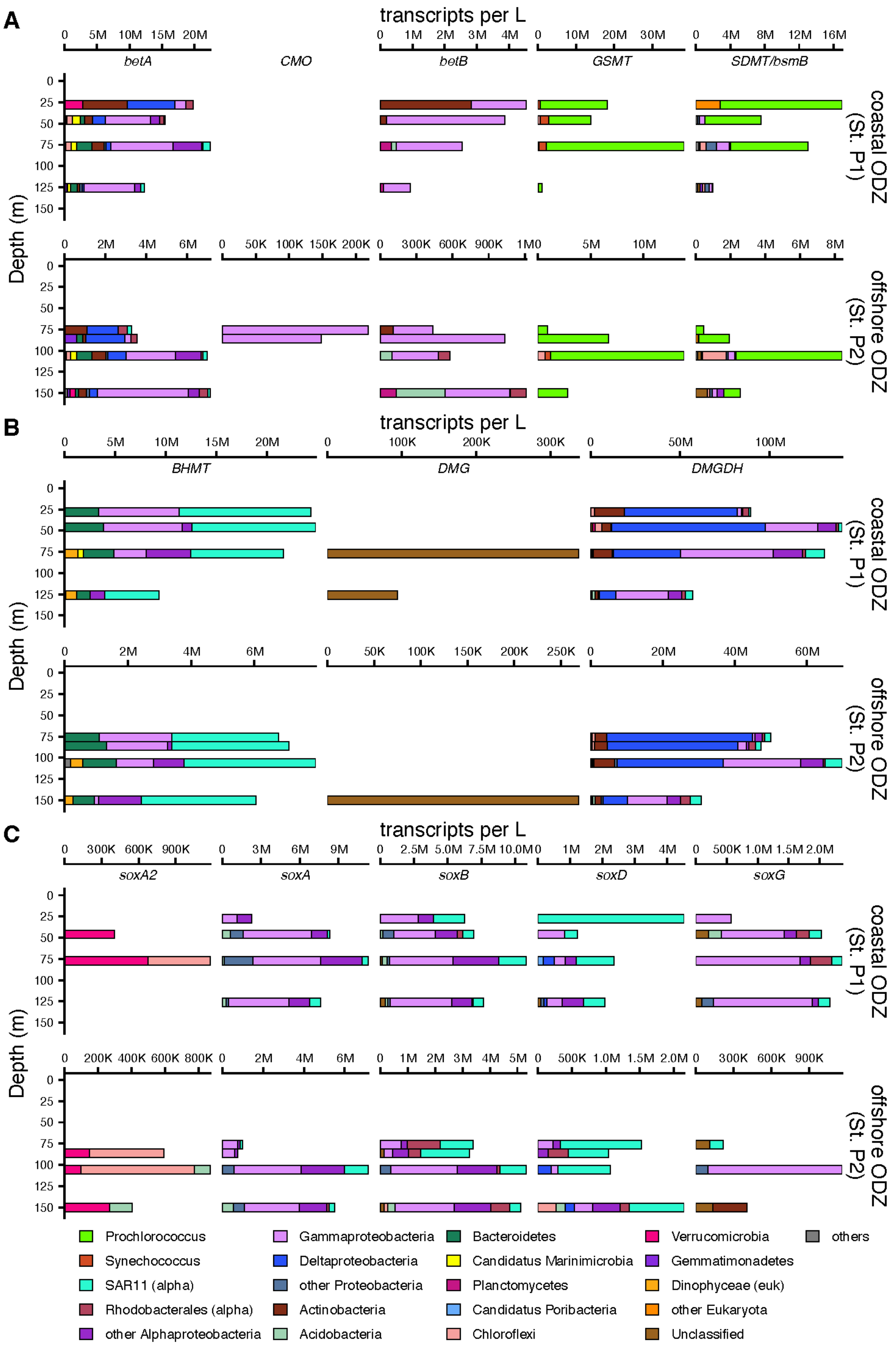


**Figure S11:** Profiles of glycine betaine (GBT) metabolic pathways expressed at the offshore ODZ station (St. P2) and coastal ODZ station (St. P1) with taxonomy. The x-axes are free for each gene. Genes: *betA*, choline dehydrogenase; *CMO*, choline monooxygenase; *betB*, betaine aldehyde dehydrogenase; *GSMT*, glycine-sarcosine methyltransferase; *bsmB/SDMT*, sarcosine dimethyltransferase; *BHMT*, betaine-homocysteine methyltransferase; *DMG*, dimethylglycine oxidase; *DMGDH*, dimethylglycine dehydrogenase; *SoxABDG*, heterotetrameric sarcosine oxidase. Pathway details and KEGG numbers for each gene provided in Table S6.

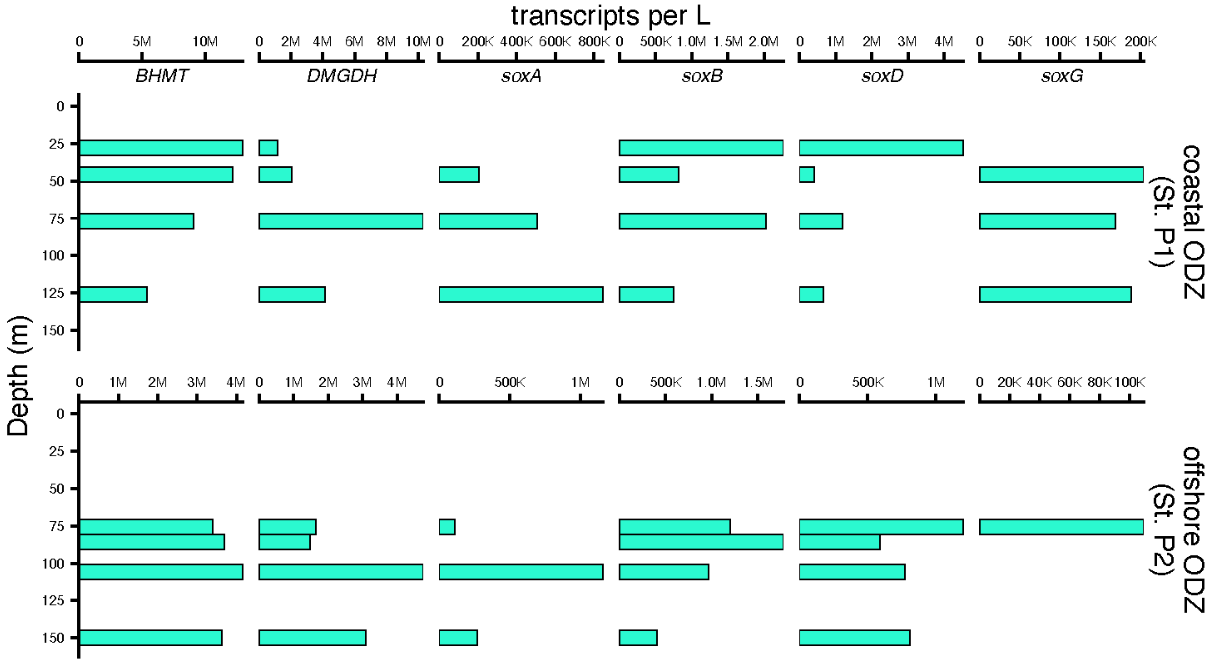


**Figure S12:** Profiles of glycine betaine (GBT) transformation by SAR11. Genes: *BHMT*, betaine-homocysteine methyltransferase; *DMGDH*, dimethylglycine dehydrogenase; *SoxABDG*, heterotetrameric sarcosine oxidase. Pathway details and KEGG numbers for each gene provided in Table S6.
